# Separable regulation of *POW1* in *TAF2*-mediated grain development and BR-mediated leaf angle formation in rice

**DOI:** 10.1101/830620

**Authors:** Li Zhang, Ruci Wang, Yueming Wang, Yufang Xu, Shuang Fang, Jinfang Chu, Shanguo Yao

## Abstract

Leaf angle is one of the key factors determining rice plant architecture. However, improvement of the leaf angle appears to be unsuccessful in practical breeding because of the simultaneous occurrence of unfavorable traits such as grain size reduction. In this study, we identified the *pow1* (*<u>p</u>ut <u>o</u>n <u>w</u>eight <u>1</u>*) mutant with enlarged grain size and leaf angle, typical brassinosteroid (BR)-related phenotypes caused by excessive cell proliferation and cell expansion. We show that *POW1* encodes a novel protein functioning in grain size regulation by repressing the transcription activity of the interacting protein TAF2, a highly conserved member of the transcription initiation complex TFIID. Loss of function of *POW1* increases the phosphorylation of OsBZR1 and decreases the inhibitory effect of OsBZR1 on the transcription of BR biosynthesis genes *OsDWARF4* (*D4*) and *D11*, thus participates in BR-mediated leaf angle regulation. The separable functions of *POW1* in grain size and leaf angle control provide a promising strategy to design high-yielding varieties in which both traits would be favorably developed, i.e., compact plant architecture and increased grain size, thus would promote the high-yield breeding a step forward in rice.

## INTRODUCTION

Rice is one of the most important food crops worldwide, and increasing grain yield remains the major challenge for most rice growing areas. The grain yield of rice is determined by three major factors, including the number of panicles per unit area, number of filled grains per panicle, and grain weight. To date, extensive attention has been paid to grain size because it is a major trait that determines grain weight and thus final yield in cereal crops, and it is one of the major targets to be selected during domestication and breeding. Grain size is specified by three components, including the length, width and thickness, and many genes or QTLs have recently been identified for their function in grain size regulation (Zuo and Li, 2014; Li et al., 2018). These regulators could be classified into multiple pathways, including mitogen-activated protein kinase signaling, ubiquitin mediated degradation, G protein signaling, phytohormone signaling, and transcriptional regulation, and all these pathways control grain size by ultimately affecting the same cellular processes of cell proliferation and/or cell expansion in the spikelet hull (Li et al., 2018). However, the molecular mechanisms underlying most of the regulators of grain size control remain largely unknown.

As another key factor that determines rice grain yield, the number of panicles per unit area is affected by the plant tillering ability and planting density, and the latter largely depends on plant architecture. Among the factors involved in determining plant architecture, BR shows great potential in improving this trait for its remarkable function in rice leaf angle regulation and grain size control (Tong and Chu, 2018). An erect leaf angle is beneficial for a higher plant density and more light capture for photosynthesis (Tian et al., 2019), which would thus result in a grain yield increase (Sakamoto et al., 2006). To date, studies on BR have exposed the double faced function of the phytohormone in plant development, i.e., compact stature brought by BR deficiency is always accompanied by small grain size, such as *d11*, *d2*, *brd1*, *dlt*, *d61-1* and *d61-2* (Yamamuro et al., 2000; Mori et al., 2002; Hong et al., 2003; Tanabe et al., 2005; Tong et al., 2009), while an enlarged grain size resulting from an excessive amount of BR often develops along with a loose stature, such as that observed for *OsBZR1*-OE, *GSK2*-RNAi, and *D11*-OE transgenic plants (Tong et al., 2012; Zhu et al., 2015). Fortunately, exceptions were observed for several cases. For example, mutation of *D4* results in plants with erect leaves without a grain size change because *D4* only contributes additional levels of bioactive BR synthesis required for normal leaf inclination and not for reproductive development (Sakamoto et al., 2006), and transgenic plants with a slightly decreased *OsBRI1* expression level exhibit a reduced leaf angle and unchanged grain size (Morinaka et al., 2006). Although grain size reduction was prevented successfully in these researches during plant architecture modification, the question arises as to whether we could raise rice plants with both traits favorably developed, i.e., a reduced leaf angle and an increased grain size.

Plant organs all achieve their final size and shape via common paths of cell division and cell expansion. Therefore, numerous reports have been dedicated to identify the way to the ultimate regulation of the genes related to these cellular processes. TAFs (TATA-box binding protein Associated Factors), as components of the TFIID complex critical for eukaryotic gene transcription (Gupta et al., 2016; Nogales et al., 2017), are reported to be essential for cell cycle progression. A TS mutation in CCG1/TAF1 in hamster cells provokes G_1_/S arrest (Sekiguchi et al., 1991), and TAF9 inactivation in chicken DT40 cells causes cell cycle arrest and apoptosis (Chen and Manley, 2000). Mouse cells that lacked TAF10 were found to be blocked in the G_1_/G_0_ phase and underwent apoptosis (Metzger et al., 1999), and a subset of TAF4b-target genes preferentially expressed in embryonic stem cells to be involved in cell cycle control (Bahat et al., 2013). Studies have also indicated that TAFs could interact with specific transcription factors (Reeves and Hahn, 2005; Garbett et al., 2007), transcription activators (Rojo Niersbach et al., 1999; Asahara et al., 2001), and components involved in epigenetic modification (Jacobson et al., 2000; Lindner et al., 2013). Although TAFs are categorized as general transcription factors, studies in *Arabidopsis* has revealed their functions in specific plant developmental processes such as light signaling (Bertrand et al., 2005), flowering time (Eom et al., 2018), pollen tube growth (Lago et al., 2005), and meristem activity and leaf development (Tamada et al., 2007). Nevertheless, more studies are required to fully expand our knowledge on TAFs in rice.

In this study, we characterized *pow1*, a recessive rice mutant with expanded grain size and enlarged leaf angle. We show that *POW1* encodes a novel protein with no transactivation activity, and functions in grain size regulation by repressing the transcriptional activity of the interactor TAF2, a highly conserved member of the TFIID complex. Loss of function of *POW1* increases the phosphorylation of OsBZR1, and decreases the inhibitory effect of OsBZR1 on the transcription of BR biosynthesis genes. Our results suggest that POW1 functions both upstream and downstream of BR signaling pathway, thus affects leaf angle formation by participating in BR homeostasis maintenance. The separable functions of *POW1* in grain size and leaf angle control provide a novel strategy to design rice plants in which both traits would be favorably developed.

## RESULTS

### Phenotypic analysis of the *pow1* mutant

By screening the NaN_3_-mutagenized M_2_ library in the background of the *japonica* rice cultivar KY131, we isolated a mutant with remarkably enlarged grain length, width and thickness compared to that of the wild type (WT, Figures 1A to 1C). The increase could reach approximately 20.0% for grain length, 23.2% for grain width, and 23.7% for grain thickness (Figure 1D), and the mutant was then designated *pow1* (*<u>p</u>ut <u>o</u>n <u>w</u>eight <u>1</u>*). Planting of the M_3_ population indicated that *pow1* displayed a loose plant architecture (Figure 1E), as featured by the near evenly extended flag leaf (Figure 1F). Detailed observation indicated that the leaf angle of *pow1* was approximately 98°, compared to approximately 34° of that of the WT. In addition, *pow1* also showed an overall expanded size of plant organs, including panicle, culm, leaf and all reproductive tissues (Supplemental Figure 1), suggesting the fundamental role of *POW1* during rice development.

**Figure 1.**
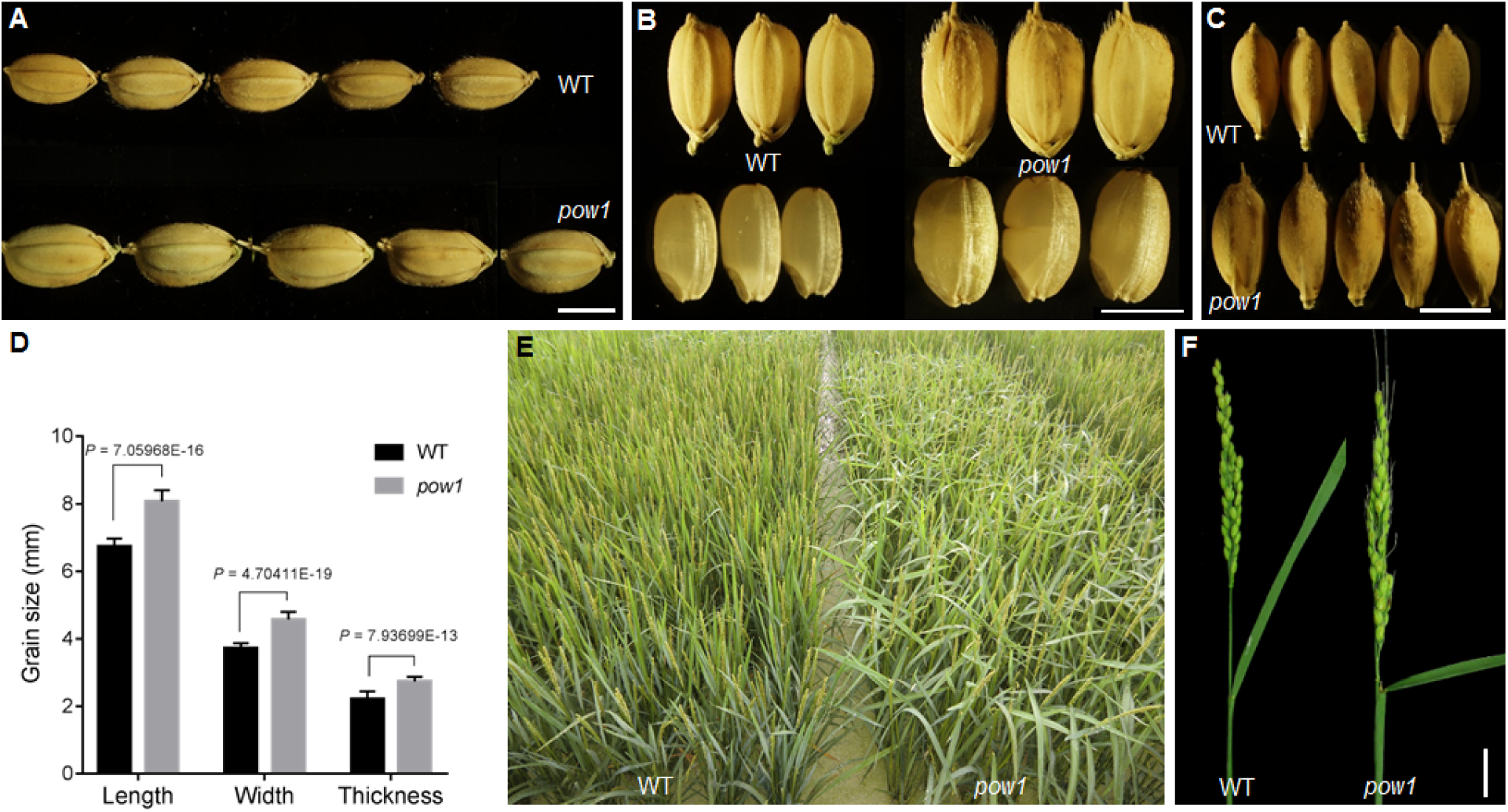
Phenotypic analysis of the *pow1* mutant. **(A)** to **(C)** Grain size comparison. Scale bars, 5 mm. **(D)** Statistical analysis of grain size. Data are means ± SE (*n* = 30). *P* values from the student’s *t*-test of *pow1* against WT were indicated. **(E)** Population morphology of *pow1* and WT. Indicating the randomly extended leaves in the mutant. **(F)** Comparison of leaf angle of the main panicle between *pow1* and WT. Scale bar, 3 cm.

Scanning Electron Microscope (SEM) observation indicated that the outer epidermal cells in the central region of the lemma in *pow1* were longer and wider than those in the WT (Figures 2A to 2C), and the increased size in the two dimensions was also confirmed by the comparison of the inner epidermal cells between *pow1* and the WT (Supplemental Figures 2A and 2B). The cell size increases in length (17.9%) and width (18.8%) were less than that of the grain size increase (Figure 1D), suggesting that *POW1* affects cell expansion, as well as cell division during grain development.

**Figure 2.**
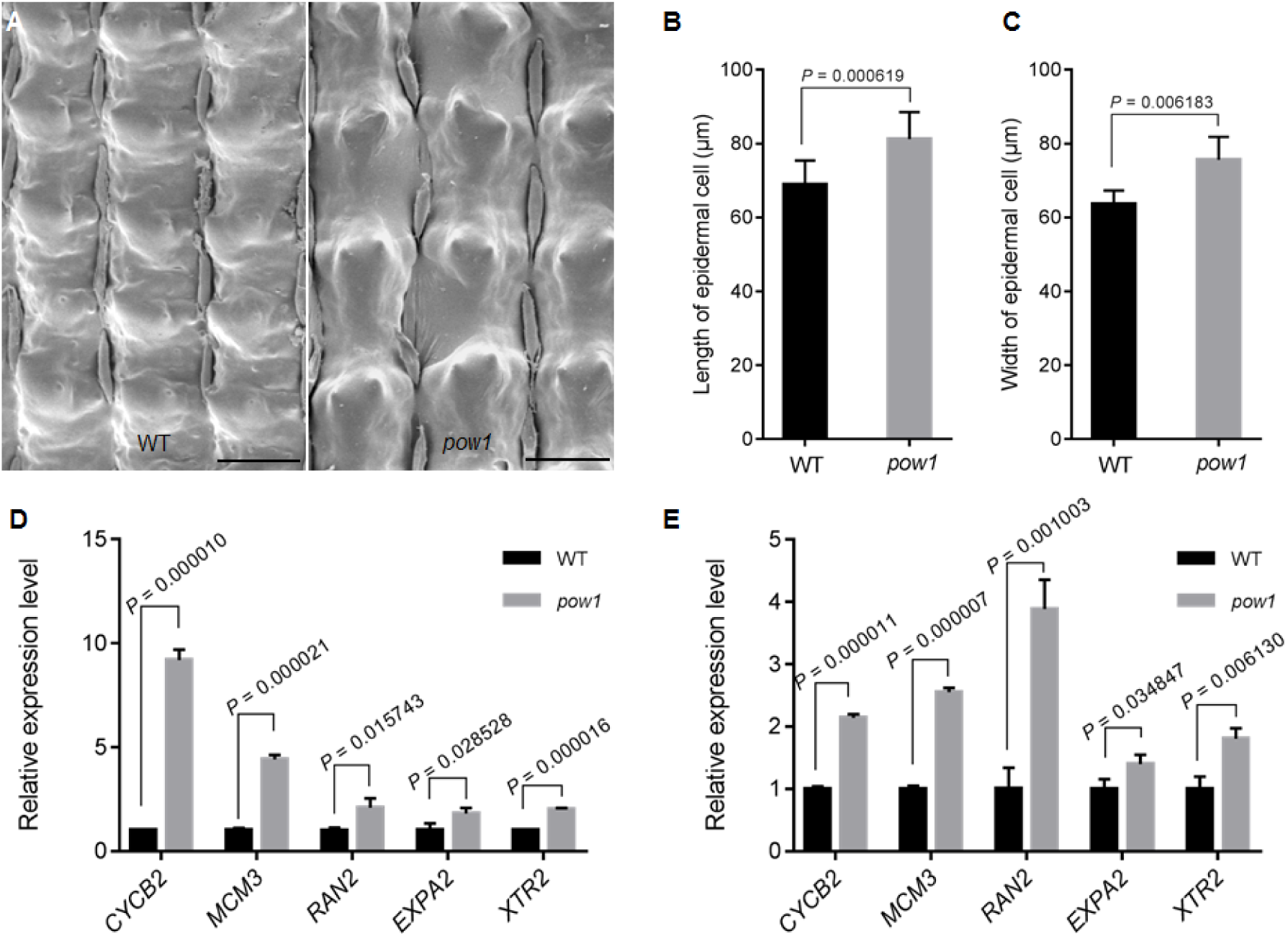
Mutation of *POW1* enhances cell division and cell expansion. **(A)** SEM observation of the outer epidermal cells in the lemma of *pow1* and WT. Scale bars, 50 μm. **(B)** and **(C)** Statistical analysis of length and width of the epidermal cells shown in **(A)**. Data are means ± SE (*n* = 100). *P* values from the student’s *t*-test of *pow1* against WT were indicated. **(D)** and **(E)** Expression analysis of cell division and cell expansion related genes in 2 mm inflorescences and lamina joints of *pow1* and WT. The transcript levels were normalized against WT, which was set to 1. Data are means ± SE (*n* = 3). *P* values from the student’s *t*-test of *pow1* against WT were indicated.

The effects of *POW1* on the cellular processes of cell division and cell expansion were further observed for the enlarged leaf angle of the mutant. We found that before the leaf angle formed, no size difference could be observed for cells in the adaxial side of the lamina joint between *pow1* and the WT, whereas the number of cell layers from the vascular bundle of xylem to the margin was clearly more in *pow1* (6∼7) than that in the WT (3∼4) (Supplemental Figure 3A), indicating the involvement of *POW1* in cell division. During the formation of the leaf angle, the number of cell layers in the adaxial side of the lamina joint increased in both *pow1* (8∼10) and the WT (6∼7) (Supplemental Figure 3B). Although cell growth was observed for both the WT and *pow1* during this stage, the increase in the cell size was much obvious in *pow1* than that in the WT (Supplemental Figures 3A and 3B), suggesting that *POW1* also functions in cell expansion.

Investigation of the expression of cell cycle and cell expansion related genes indicated that in both young inflorescence and lamina joint, *pow1* exhibited much higher expression levels of the cell division marker genes, such as *MCM3*, *CYCB2*, and *RAN2*, and the cell expansion marker genes, such as *XTR2* and *EXPA2* (Duan et al., 2012; Jiang et al., 2012) (Figures 2D and 2E). These results are well consistent with the phenotypic observations that *POW1* functions in grain size and leaf angle development by ultimately affecting both cell division and cell expansion.

### *POW1* is ubiquitously expressed and encodes a protein with unknown function

To understand the molecular function of *POW1*, we cloned the causal gene by map-based cloning. After sequencing of the fine-mapped region, we found a single mutation from G to T occurred in *pow1* in an annotated gene *Os07g07880* (http://rice.plantbiology.msu.edu/), which caused an amino acid change from Arginine to Serine (Figure 3A).

**Figure 3.**
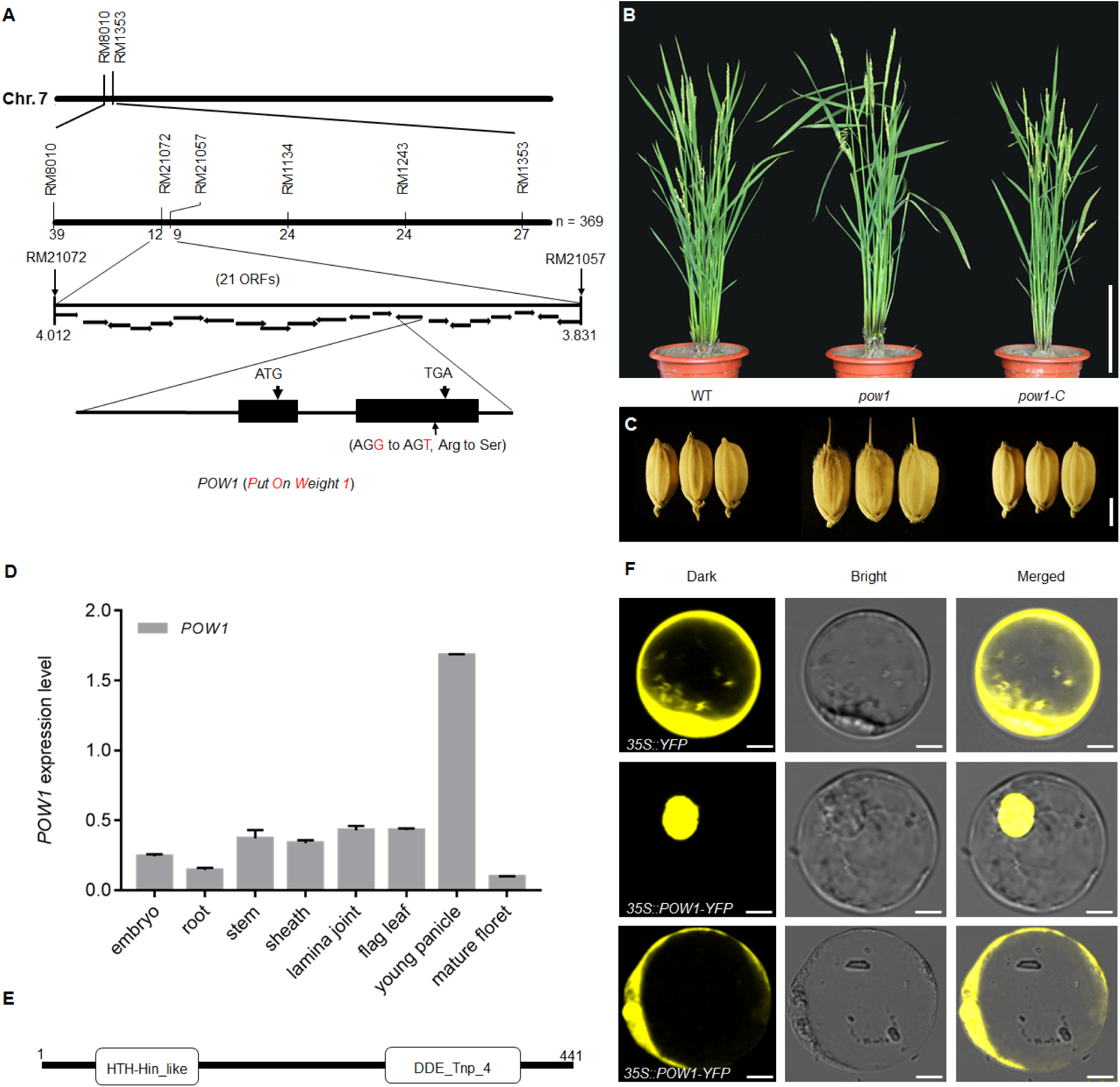
Map-based cloning of *POW1*. **(A)** Identification of the *POW1* candidate. *POW1* is located between RM21072 and RM21057 on chromosome 7. Sequence comparison found one mutation from G to T occurred in the mutant, which results in an amino acid change from Arg to Ser. **(B)** and **(C)** Phenotypes of whole plants and grains of *pow1*, WT and complemented T_0_ line (*pow1*-C). Scale bars, 20 cm for **(B)** and 5 mm for **(C)**. **(D)** Expression analysis of *POW1* in various plant tissues. Data are means ± SE (*n* = 3). **(E)** Protein structure of POW1. **(F)** Subcellular localization of POW1. POW1 is localized in the nucleus in most cells. For a small portion of cells (≈ 10%), however, POW1-YFP is observed in cytoplasm and membrane, as well as in nucleus. Scale bars, 10 μm.

To verify that *Os07g07880* is actually *POW1*, genomic DNA, including 2.2-kb upstream of ATG and 948-bp downstream of TGA, was amplified from the WT and transformed into *pow1.* We found that all 38 T_0_ transgenic plants showed overall phenotypes, including grain size and leaf angle, resembling that of the WT (Figures 3B and 3C), demonstrating that the *pow1* phenotypes are caused by the point mutation in *Os07g07880*.

*POW1* is ubiquitously expressed in all tissues, including the root, embryo, stem, inflorescence, flag leaf and lamina joint (Figure 3D), which fits well with the expanded size of all organs in the mutant (Figure 1 and Supplemental Figure 1). The highest expression of *POW1* was observed in young panicles, and the expression level of the gene decreased during panicle maturation (Figure 3D). This result suggests that *POW1* plays a critical role in early inflorescence development.

*POW1* is predicted to encode a homeodomain-like protein with a putative Helix-Turn-Helix DNA binding structure and a DDE domain (Figure 3E, Yuan and Wessler, 2011). Although POW1 appeared to belong to the superfamily of Harbinger DNA transposons that possess nuclease activity (Kapitonov and Jurka, 1999), the typical catalytic DDE domain was disturbed in POW1 (Supplemental Figure 4). This observation suggests that POW1 might lose the activity as a transposase (Kapitonov and Jurka, 2004) and obtain a new function during the evolution, such as that reported for the *Arabidopsis* homolog *ALP1* (Kapitonov and Jurka, 2004; Liang et al., 2015).

In most of the cells (≈ 90%), POW1 is exclusively observed in the nucleus. However, for a small portion of cells (≈ 10%), we found that POW1 is also localized in the cytoplasm and membrane, as well as the nucleus (Figure 3F). Because POW1 contained a putative DNA binding domain and mostly localized in the nucleus, we subsequently questioned whether POW1 functions as a transcription factor. However, we could not detect any transactivation activity of POW1 in yeast (Supplemental Figure 5A). We further performed the transcription activity assay in rice protoplast using the luciferase coding gene as a reporter. When we transformed the GAL4-BD-POW1 fusion protein into rice protoplast, we only observed a 1.5-fold change in the luciferase expression level compared to that of the negative control (Supplemental Figure 5B). In conclusion, these results suggest that POW1 is neither a transcriptional activator nor a repressor.

### *POW1* affects leaf angle formation via BR pathway

The increase in leaf angle is the typical phenotype displayed by plants with an excessive amount of BR. In particular, the loose population structure of *pow1* highly resembled plants that ectopically express *At-CYP90B1* or *Zm-CYP* (Wu et al., 2008), which are homologs of rice *D4* and *D11*, respectively. Therefore, we first quantified the endogenous BR content in the mutant. We found that the content of castasterone, a likely end product of the BR biosynthetic pathway in rice(Kim et al., 2008), was obviously higher in *pow1* than that in the WT (Figure 4A). Consistent with the higher endogenous BR content, we found that the expression levels of BR biosynthesis genes in the mutant were enhanced for *D4* in the flag leaves and *D11* in the young inflorescence, while the expressions of the other two genes *BRD1* and *D2* were not significantly changed between *pow1* and the WT (Figure 4B). The elevated BR level in the mutant was further confirmed by seedling treatment with Brassinazole (BRZ), a BR biosynthesis inhibitor that specifically blocks BR biosynthesis by inhibiting the cytochrome P450 steroid C-22 hydroxylase encoded by the *DWF4* gene (Asami et al., 2001). We found that the position of the second lamina joint in *pow1* was obviously higher than that in the WT under control conditions, and the phenotype of *pow1* seedlings treated with 5 μM BRZ highly mimicked that of the WT without BRZ treatment (Figure 4C). Taken together, these results indicate that *pow1* is a BR-overproducing mutant.

**Figure 4.**
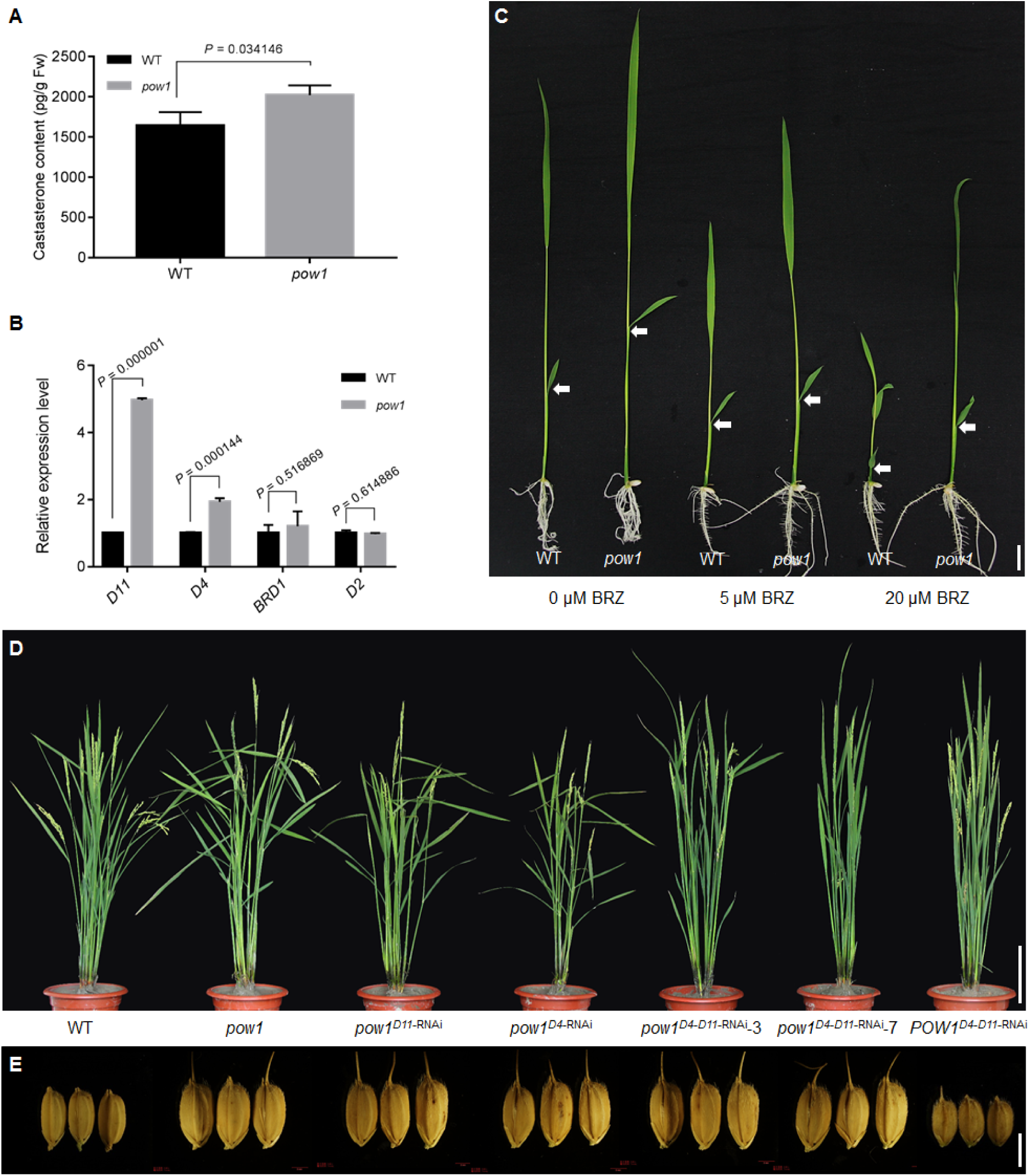
*pow1* is a BR-overproducing mutant. **(A)** Endogenous BR quantification. Two-month-old field grown plants were sampled. Data are means ± SE (*n* = 3). *P* values from the student’s *t*-test of *pow1* against WT were indicated. **(B)** Expression analysis of BR biosynthesis genes in 2 mm inflorescences for *D11* and flag leaves for *D4*. The transcript levels were normalized against WT, which was set to 1. Data are means ± SE (*n* = 3). *P* values from the student’s *t*-test of *pow1* against WT were indicated. **(C)** BRZ treatment. The sterilized seeds were sowed on half strength MS medium that contained different concentrations of BRZ and cultured for one week after germination. The white arrows indicate the second lamina joints. Scale bar, 1 cm. **(D)** and **(E)** Comparison of leaf angle and grain size among the *D4* and *D11* RNAi transgenic lines under the background of *pow1* and WT. Scale bars, 20 cm for **(D)** and 5 mm for **(E)**.

To verify whether the mutant phenotype was caused by ectopic expression of the BR biosynthesis genes *D4* and *D11*, we constructed RNAi transgenic plants of the two genes under *pow1* background. We could not observe phenotypic recovery in the *D4* or *D11* single RNAi transgenic plants (Figures 4D and 4E), which might be due to the higher expression level of the other gene (Supplemental Figure 6A). When simultaneously knocking down these two genes (Supplemental Figure 6A), we found that the *pow1^D4^*^−*D11*-RNAi^-3 plant exhibits a leaf angle much smaller than *pow1*, and the leaf angle of *pow1^D4^*^−*D11*-RNAi^-7 is comparable to that of *D4* and *D11* double RNAi transgenic plants under the WT background (*POW1^D4^*^−*D11*-RNAi^; Figure 4D), which suggests that *POW1* controls leaf angle formation by regulating the transcription of *D4* and *D11*. Unexpectedly, we could not observe remarkable changes in grain size in all double RNAi lines under the *pow1* background, although the grain size of the *POW1^D4^*^−*D11*-RNAi^ plant is sharply reduced compared to that of the WT (Figure 4E and Supplemental Figure 6B).

In addition to BR biosynthesis, altered BR signaling could also affect grain size development (Yamamuro et al., 2000; Zhu et al., 2015). Our BRZ treatment indicated that *pow1* showed alleviated inhibition under 20 μM BRZ conditions (Figure 4C), suggesting that *POW1* might be involved in the BR signaling pathway. We then performed a lamina joint bending assay and found that the lamina joint bending angle in *pow1* was larger than that in the WT under all brassinolide (BL) concentrations (Figure 5A and Supplemental Figure 7A). Because coleoptile growth under BR treatment is an important BR responsive phenotype (Yamamuro et al., 2000), we further detected the coleoptile length of *pow1* and WT seedlings under a series of BL concentrations. No clear difference could be observed for the growth rate of coleoptile between *pow1* and the WT under the low BL concentrations; however, *pow1* displayed a 36% more increase in the coleoptile length than in the WT under 5 μM BL treatment (Figure 5B and Supplemental Figure 7B). These results suggest that *POW1* is involved in BR signaling, as well as BR biosynthesis.

**Figure 5.**
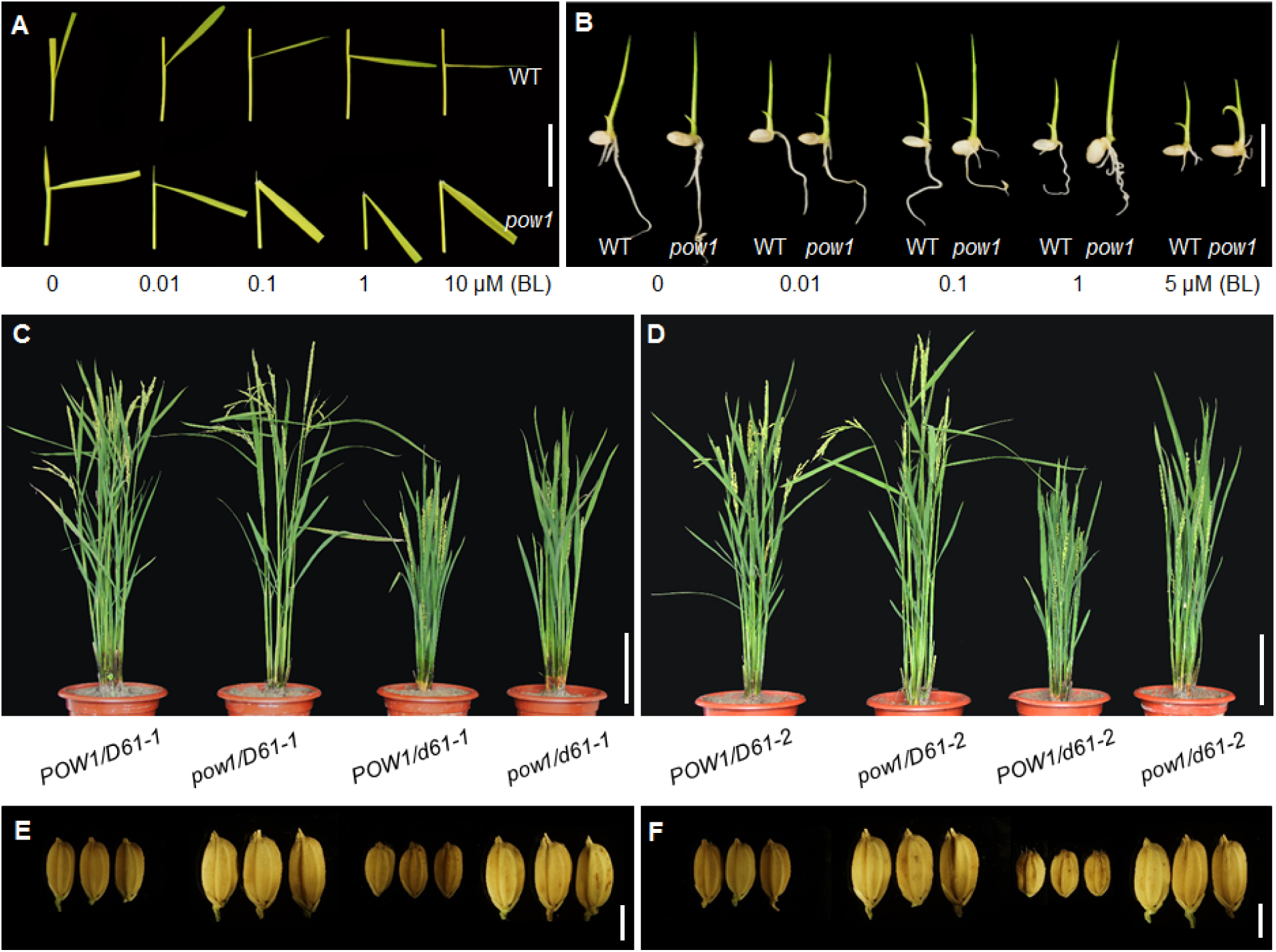
*pow1* is a BR-signaling mutant. **(A)** Lamina joint assay. The second leaf laminas of 7-day-old dark grown seedlings were excised and submerged in distilled water of different BL concentrations for 3 days. Scale bar, 1 cm. **(B)** Coleoptile elongation assay. The coleoptile length of *pow1* and WT grown for 3 days under different BL concentrations was compared. Scale bar, 1 cm. **(C)** to **(F)** Comparison of leaf angle and grain size between *pow1* and *d61/pow1* double mutants. Two allelic mutants of *d61-1* and *d61-2* were used for analysis. Scale bars, 20 cm for **(C)** and **(D)**, and 5 mm for **(E)** and **(F)**.

Consistent with the hypersensitive response of *pow1* to BR, we found that expression of the genes positively involved in BR signaling, including *OsBRI1*, *RAVL1*, *ILI1* and *OsBZR1*, was all significantly induced under the mutant background (Supplemental Figure 7C). Therefore, we attempted to understand whether BR signaling participates in *POW1*-mediated grain size regulation by crossing *pow1* with *d61-1* and *d61-2*, two allelic mutants of the BR receptor gene *OsBRI1* that show less sensitivity to BL than their relative WT (Yamamuro et al., 2000). To exclude the background noise, we selected double mutants and single mutants from the same genetic population for phenotypic comparison. We found that the leaf angle of the double mutant *pow1/d61-2* is completely recovered to that of the *POW1/d61-2* single mutant, while the leaf angle in *pow1/d61-1* is only partially rescued compared to that in *POW1/d61-1* (Figures 5C and 5D; Supplemental Figures 8A and 8B), which should be because *d61-1* is a weak allele (Yamamuro et al., 2000). Surprisingly, we could not yet observe any difference in the grain size among *pow1/D61-1*, *pow1/D61-2* and the double mutants (Figures 5E and 5F; Supplemental Figures 8C and 8D). These results suggest that POW1 regulates grain size downstream of BR signaling.

### POW1 interacts with and regulates grain size through TAF2

The BR biosynthesis genes *D4* and *D11* show altered transcription under the *pow1* background (Figure 4B), whereas POW1 does not appear to function as a transcription factor (Supplemental Figures 5A and 5B). We first speculated that POW1 might act as an epigenetic regulator like its *Arabidopsis* homolog ALP1 (Liang et al., 2015). However, this possibility was excluded because no significant alteration in the H3K27me3 levels was observed within either *D4* or *D11* gene locus (Supplemental Figure 5C). Therefore, we queried whether POW1 carries out transcriptional regulation by associating with other factors (Sridhar et al., 2006; Jiang et al., 2018). To this end, we screened its potential interactors using a cDNA library with the yeast two-hybrid system. Among the 274 sequenced positive colonies, we were highly interested in one POW1-interacting insert (Figures 6A to 6C), which is the C-terminus of *Os09g24440* (http://rice.plantbiology.msu.edu/) predicted to encode the transcription initiation factor TFIID subunit 2 (TAF2, https://www.ncbi.nlm.nih.gov/nucleotide/XM_015756386.2). As a highly conserved member, TAF2 plays a crucial role in cell cycle progression (Lago et al., 2004), which is closely related to the function of *POW1* (Figures 2D and 2E). Therefore, we speculated that these two factors might work as a complex to mediate rice development. Because *TAF2* has no paralogs in the rice genome (https://blast.ncbi.nlm.nih.gov/Blast), and knockout of the core components, such as *TAF1* or *TAF6*, typically causes a lethal phenotype (Lago et al., 2005; Waterworth et al., 2015), we constructed *TAF2*-RNAi transgenic plants under the WT background to study its function. Among the 52 independent transgenic lines, we selected three lines that showed differentially repressed expression of *TAF2* for phenotypic analysis. We found that all the three lines exhibited phenotypes contrary to that of *pow1*, including a reduced grain size, decreased leaf angle, and shortened plant height compared to those of WT (Figures 7A to 7C), and the grain size reduction in the *TAF2*-RNAi plants was due to the decrease in both cell size and cell number (Supplemental Figure 9). Consistently, we found that expression of the cell division markers, such as *MCM3* and *CYCB2*, and the cell expansion gene *EXPA2*, was all repressed along with the downregulation of *TAF2* (Figure 7D), which is in sharp contrast to the upregulation of these genes under the *pow1* background (Figures 2D and 2E). We further observed that as a predicted transcription initiation factor, TAF2 could activate significantly the expression of all the three marker genes (Figure 7E and Supplemental Figure 10A), which explained well the differential expression of these genes between WT and *TAF2*-RNAi transgenic plants (Figure 7D). Furthermore, we found that the activation activity of TAF2 on *CYCB2* could be significantly repressed by POW1, while the mutated protein (mPOW1) showed opposite effect on *CYCB2* expression (Figure 7F and Supplemental Figure 10B). By contrast, there was no significant difference between POW1 and mPOW1 in the activation of *MCM3* and *EXPA2* by TAF2 (Figure 7F and Supplemental Figure 10B), indicating that the upregulated expression of these genes in *pow1* is not due to the differential effects of POW1 and mPOW1 on TAF2 activation activity. Because the expression level of *TAF2* was obviously higher in *pow1* than that in WT (Figure 6D), the upregulated expression of *MCM3* and *EXPA2* in the mutant should be due to the more accumulated transcription activator TAF2. Totally, these observations indicate that POW1 antagonizes TAF2 in regulating downstream gene transcription, and the lifted repression on the transactivation activity of TAF2 by mPOW1 well explained that in the inflorescence with the highest *POW1* expression (Figure 3D), the transcription of *CYCB2* was much enhanced than that of all the other genes under the *pow1* background (Figure 2D).

**Figure 6.**
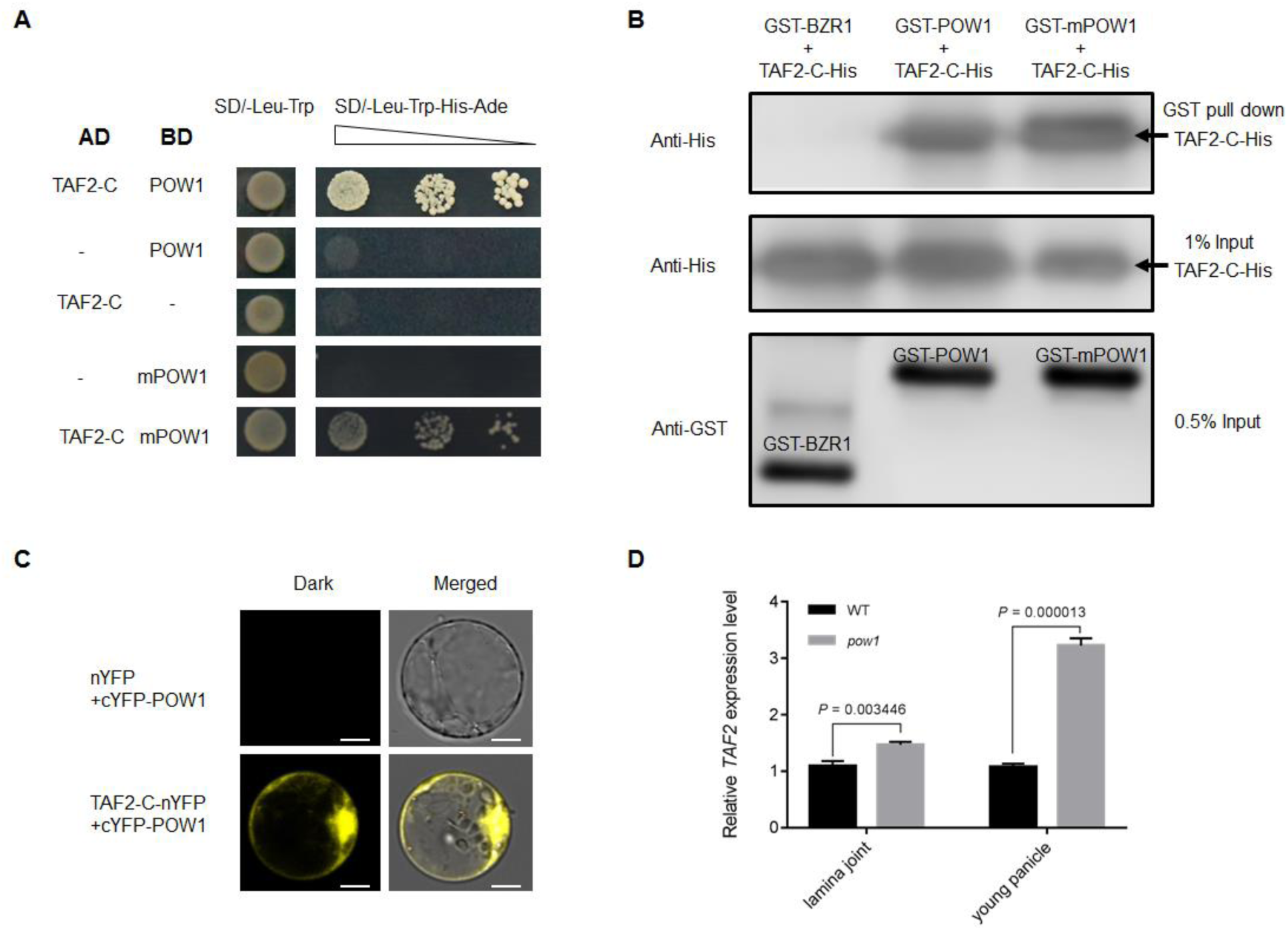
Transactivation assay of TAF2 and its interaction with POW1. **(A)** to **(C)** Interaction assay between POW1/mPOW1 and TAF2. GST-OsBZR1 was used as negative control. Scale bars, 10 μm. **(D)** Expression analysis of *TAF2* in the lamina joint and young panicle of *pow1* and WT. The transcript levels were normalized against WT, which was set to 1. Data are means ± SE (*n* = 3). *P* values from the student’s *t*-test of *pow1* against WT were indicated.

**Figure 7.**
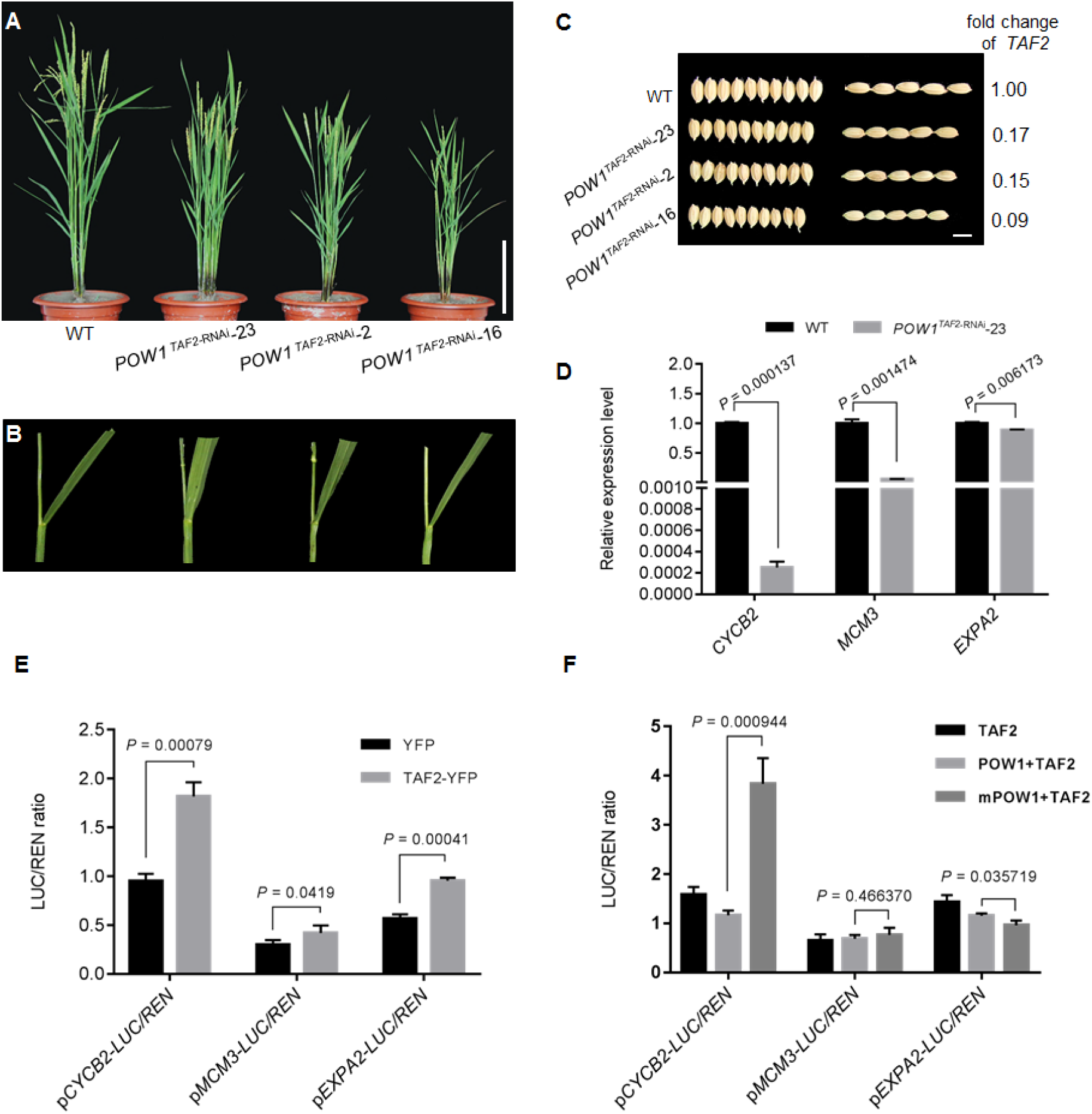
*TAF2* is a positive regulator for grain size. **(A)** to **(C)** Phenotypes of plants, leaf angle, and grain size in the *TAF2*-RNAi transgenic lines. Scale bars, 20 cm for **(A)** and 5 mm for **(C)**. **(D)** Expression analysis of cell division and cell expansion related genes in the *TAF2*-RNAi transgenic line. The transcript levels were normalized against WT, which was set to 1. Data are means ± SE (*n* = 3). *P* values from the student’s *t*-test of the *TAF2*-RNAi transgenic line against WT were indicated. **(E)** TAF2 could effectively activate the transcription of cell division and cell expansion genes. Data are means ± SE (*n* = 3). *P* values calculated from the student’s *t*-test were indicated. **(F)** Activation of TAF2 on the transcription of cell division and cell expansion genes were differentially affected by POW1 and mPOW1. Data are means ± SE (*n* = 3). *P* values from the student’s *t*-test of POW1 against mPOW1 were indicated.

Very interestingly, we found that although the leaf angle of *POW1^TAF2^*^−RNAi^ transgenic plants appeared to be smaller than that of WT, the change of leaf angle did not seem to be proportional to the expression level of *TAF2* (Figures 7B and 7C). Therefore, we further explored the function of *TAF2* in POW1-mediated leaf angle regulation by creating the *TAF2*-RNAi transgenic plants under the mutant background (Figures 8A and 8D). We found that knockdown of *TAF2* could largely recover the grain size of *pow1* to that of the WT (Figures 8B and 8F). However, the enlarged leaf angle phenotype remained unchanged in these transgenic plants (Figure 8C), and longitudinal sections of the lamina joint of *TAF2*-RNAi transgenic plants showed that downregulation of *TAF2* did not affect cell elongation during leaf angle formation (Supplemental Figure 11), although significant effect of this gene on cell elongation was observed during grain development (Supplemental Figure 9). These results suggest that *TAF2* is not involved in POW1-mediated leaf angle formation. This conclusion was further supported by analyzing the expression levels of *D4* and *D11* in these transgenic plants because the large leaf angle of *pow1* is due to the upregulation of these two genes (Figure 4D). We found that in the *pow1^TAF2^*^−RNAi^ plants, the expression levels of these two genes were substantially decreased compared with that in *pow1* but significantly higher than that in the WT (Figure 8E), which might explain the unchanged leaf angle phenotype in these plants. Therefore, we constructed transgenic plants with simultaneously downregulated expression of *TAF2*, *D4* and *D11* under the mutant background, and the triple RNAi transgenic plants *pow1^TAF2-D11-D4^*^−RNAi^ displayed largely rescued phenotypes similar to that of the WT, i.e., decreased grain size and erect leaf angle (Figure 8). These results further demonstrate that *POW1* controls leaf angle formation via the BR pathway and regulates grain size development under the assistance of TAF2.

**Figure 8.**
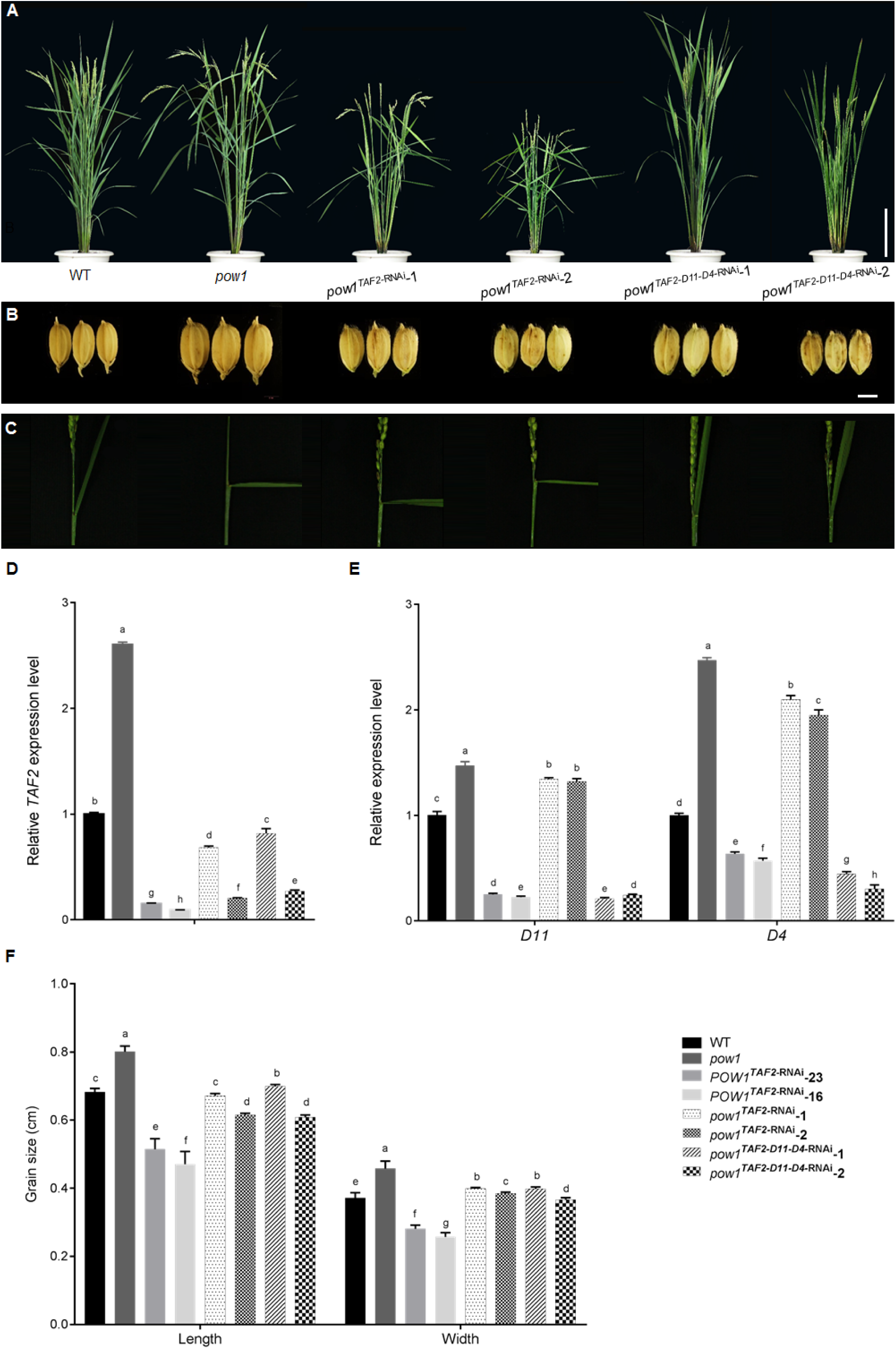
Downregulation of *TAF2* could rescue the grain size of *pow1*. **(A)** to **(C)** Phenotypes of plants, grain size, and leaf angle in the *TAF2*-RNAi transgenic lines under the *pow1* mutant background. Scale bars, 20 cm for **(A)** and 5 mm for **(B)**. **(D)** and **(E)** Expression analysis of *TAF2*, *D4* and *D11* in the RNAi transgenic lines under the background of *pow1* and WT. The transcript levels were normalized against WT, which was set to 1. Data are means ± SE (*n* = 3). Bars followed by the different letters represent significant difference at 5%. **(F)** Statistical analysis of grain length and grain width in the RNAi transgenic lines under the background of *pow1* and WT. Data are means ± SE (*n* = 15). Bars followed by the different letters represent significant difference at 5%.

### *POW1* is negatively involved in BR homeostasis possibly via GL7

Our genetic analysis clearly showed that *POW1* regulates leaf angle formation by affecting the transcription of the BR biosynthesis genes *D4* and *D11* (Figure 4D). To explore how POW1 affects the transcription of *D4* and *D11*, we focused on OsBZR1, a central transcription factor reported to mediate the negative feedback regulation of the BR biosynthesis genes by binding directly to the promoters of *D4* and *D11* (Bai et al., 2007; Qiao et al., 2017). Although we could not detect any interaction between POW1 and OsBZR1 (Supplemental Figure 12), we found that OsBZR1 showed more cytoplasm retention in *pow1* than that in WT (Figures 9A and 9B). This is quite interesting because previous studies have proven that BL induction could induce the nuclear localization of OsBZR1 (Bai et al., 2007), in contrast to the higher BR content and more cytoplasm retention of OsBZR1 under the *pow1* background. Because the subcellular localization of OsBZR1 fully reflects the phosphorylation status of the protein (Wang et al., 2012), we therefore analyzed the phosphorylation of OsBZR1 in *pow1* and WT. We found that *pow1* possessed more phosphorylated OsBZR1 than WT (Figure 9C), which is consistent well with the cytoplasm retention of OsBZR1 in the mutant. Furthermore, we also observed that the inhibitory effect of OsBZR1 on the expression of *D4* and *D11* is obviously reduced in *pow1* compared to that in WT (Figure 9D and Supplemental Figure10C), which should be due to the less nuclear localized OsBZR1 protein. Moreover, we found that similar to that observed in the BR-deficient mutants (Hong et al., 2003), the expression of *POW1* was highly enhanced in the *pow1* mutant (Figure 9E), which may represent a self-compensation effect for the mutation. And, application of exogenous BL could effectively induce the expression of *POW1* (Figure 9F). Taken together, these results suggest that POW1 is a key regulator of OsBZR1-mediated BR negative feedback loop, and is important for the stable nuclear localization of OsBZR1 when the endogenous BR content is high.

**Figure 9.**
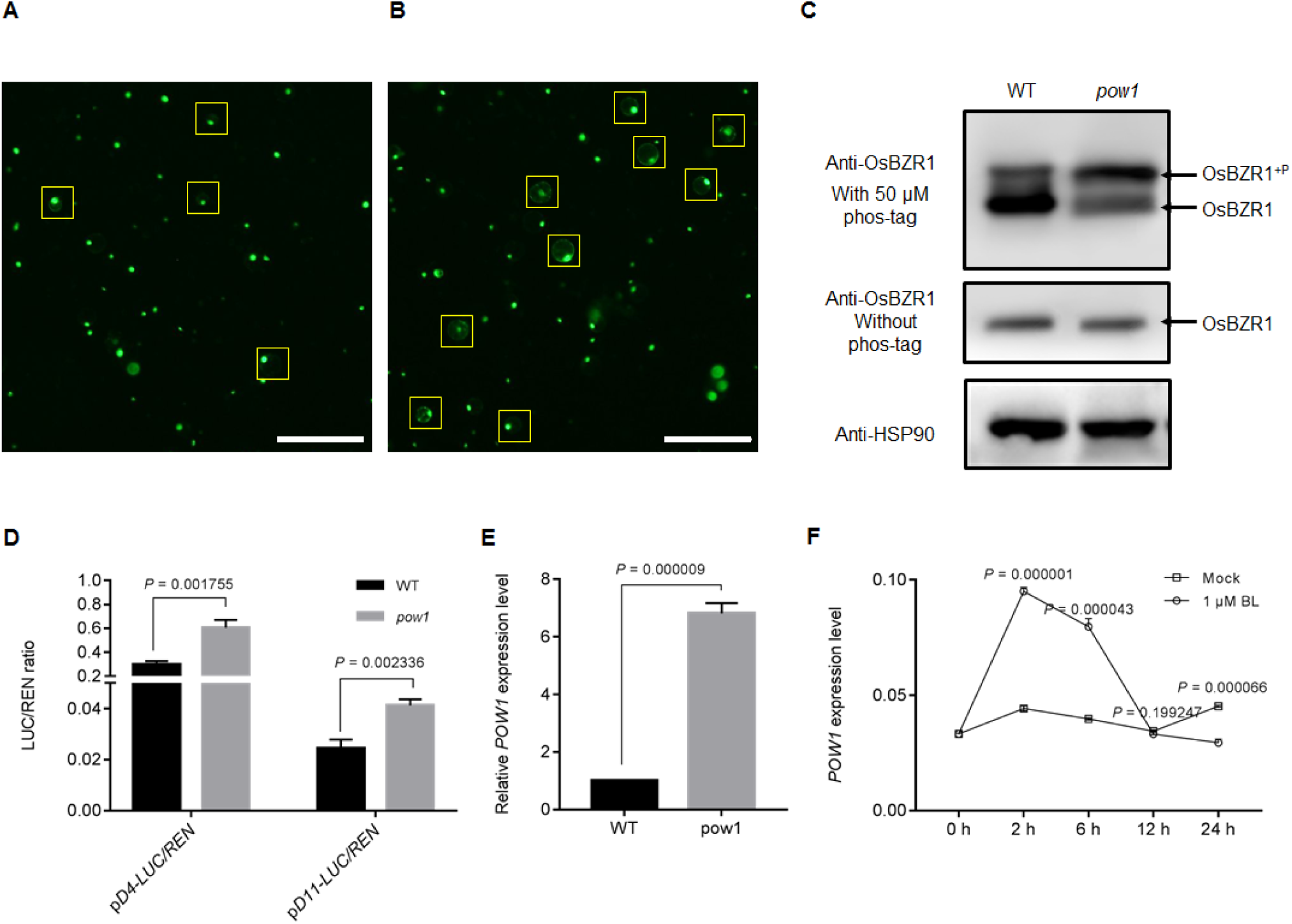
*POW1* affects BR homeostasis by altering OsBZR1 subcellular localization. **(A)** and **(B)** Protein localization. OsBZR1-YFP localizes mostly in the nuclear of WT protoplast **(A)**, while the signals were mostly observed in the cytoplasm of *pow1* protoplast **(B)**. Scale bars, 200 μm. **(C)** Phosphorylation assay. Indicating that *pow1* possesses more phosphorylated OsBZR1. **(D)** Transactivation activity assay. Indicating that the mutation of *POW1* reduces the inhibition of OsBZR1 on *D4* and *D11* expression. **(E)** Comparison of *POW1* expression between *pow1* and WT. The transcript levels were normalized against WT, which was set to 1. **(F)** BR induction assay. The whole shoots of two-week-old seedlings under 1 μM BL were collected after 0, 2, 6, 12, and 24 h treatment. Data in **(D)**-**(F)** are means ± SE (*n* = 3), and *P* values from the student’s *t*-test were indicated.

Although we revealed that POW1 controls the expression of *D4* and *D11* by affecting OsBZR1 phosphorylation, we could not detect a direct interaction between these two proteins (Supplemental Figure12). We hypothesized that OsBZR1 might regulate transcription by interacting with transcription initiation factors such as TAF2. To our regret, no interaction was detected between OsBZR1 and TAF2 (Figure 6B and Supplemental Figure 12). Therefore, we tried to screen the potential linkers of POW1 and OsBZR1 using the yeast two-hybrid system with POW1 as the bait, and identified the POW1-interacting protein GL7 (GW7, Figure 10), a major QTL involved in grain size regulation (Wang et al., 2015a; Wang et al., 2015b). Although GL7 did not interact directly with OsBZR1, it showed strong interaction with GSK2 (Figure 10), a kinase responsible for the phosphorylation of OsBZR1 (Liu et al., 2017). These observations suggest that GL7 might be the connexon in the regulation of OsBZR1 phosphorylation by POW1. However, the detailed regulatory mechanism needs to be further studied.

**Figure 10.**
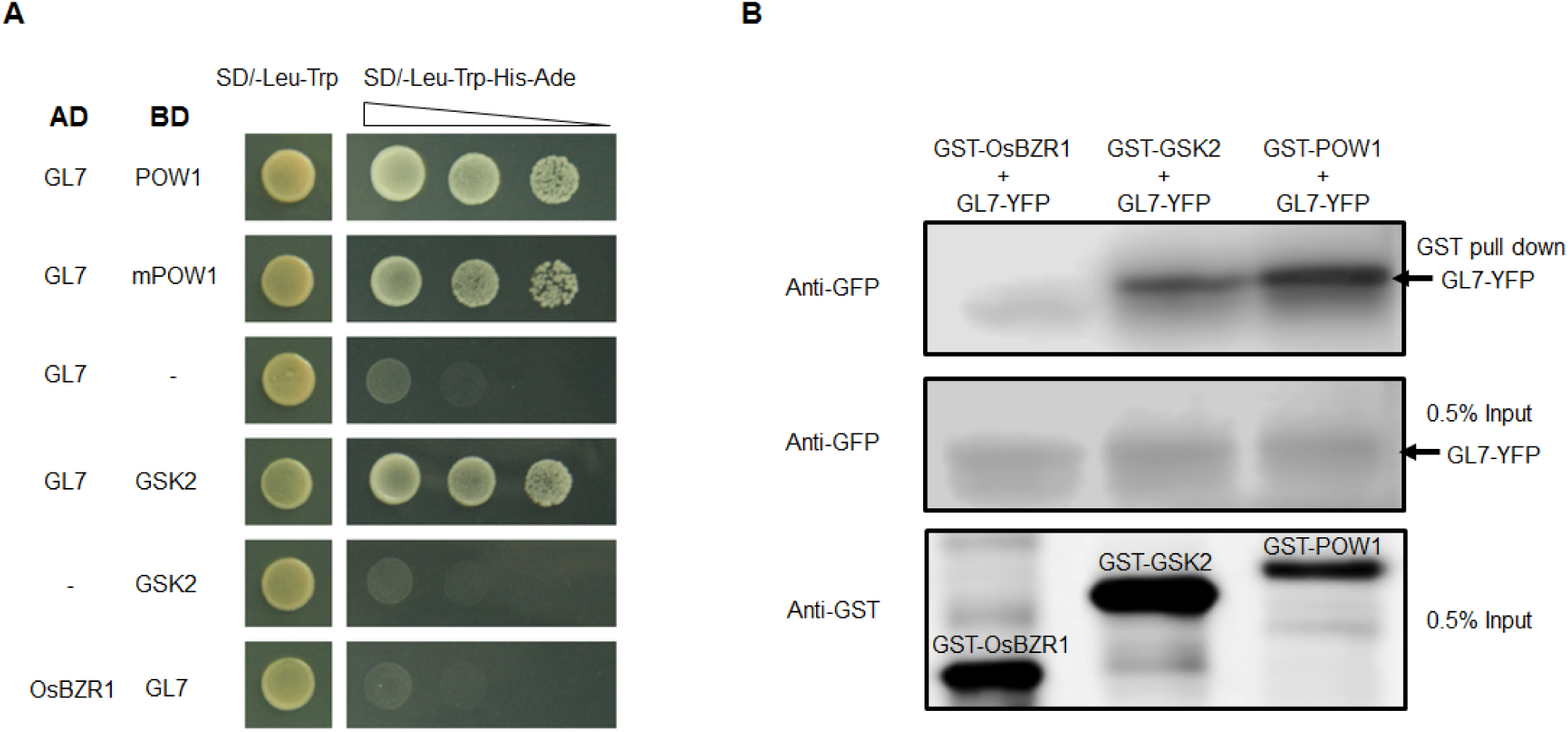
GL7 might be a linker between POW1 and BR pathway. **(A)** Interaction assay. GL7 could interact with POW1 and GSK2 but not OsBZR1 in yeast. **(B)** Semi-pull down assay. Indicating the interaction between GL7 and POW1/GSK2. GST-OsBZR1 was used as the negative control.

## DISCUSSION

Although *pow1* shows typical BR-related phenotypes of enlarged grain size and leaf angle, we provided evidence that *POW1* controls these two traits through separable routes of BR pathway and the transcription initiation factor *TAF2*, respectively. The separable functions of *POW1* in grain size and leaf angle regulation imply that POW1 occupies a critical position in integrating these two important biological processes.

### POW1 acts both upstream and downstream of the BR pathway

Loss of function of *POW1* enhanced the expression of the BR biosynthesis genes *D4* and *D11*, resulting in a higher BR content which could induce the expression of *POW1*. These results suggest that *POW1* plays a critical role in the feedback regulation of BR homeostasis, which is similar to that reported for *OsBZR1* (Bai et al., 2007). OsBZR1 is the key transcriptional regulator of the BR pathway which binds directly to the promoters of downstream targets (Tong et al., 2012; Zhu et al., 2015; Qiao et al., 2017). However, the initiation of gene transcription depends on not only specific transcription factors but also transcription initiation complex, which is consist of TFIIA, TFIIB, TFIID, TFIIE, TFIIF, TFIIH and RNA polymerase II (Gupta et al., 2016). So far, knowledge on the function of TAFs during rice development remains blank, and our findings show clearly that TAF2 plays a pivotal role in cell division and cell expansion. As the core TFIID component, TAF2 is always first recruited to the transcription initiation region (Nogales et al., 2017), thus enables the subsequent package of the mature transcription initiation complex. However, for the transcription of some inducible genes, other additional factors also participate in this basic transcription initiation process (Weake and Workman, 2010). As an important growth regulator, BR could affect rice development by inducing the expression of many genes (Tong and Chu, 2018), and OsBZR1 thus might act as an additional factor to join the *TAF2/POW1*-mediated transcription of downstream genes, including those involved in cell division and cell elongation. Because POW1 interacts with and affects the transactivation activity of TAF2, and participates in BR homeostasis through impacting on OsBZR1 phosphorylation, the BR-inducible *POW1* thus functions possibly both upstream and downstream of *OsBZR1*-mediated BR signaling. We propose that *POW1* regulates the transcription of downstream genes via a fine equilibration among OsBZR1 (BR content), TAF2 and POW1, while the equilibrium point might be different for leaf angle formation and grain size determination (Figure 11).

**Figure 11.**
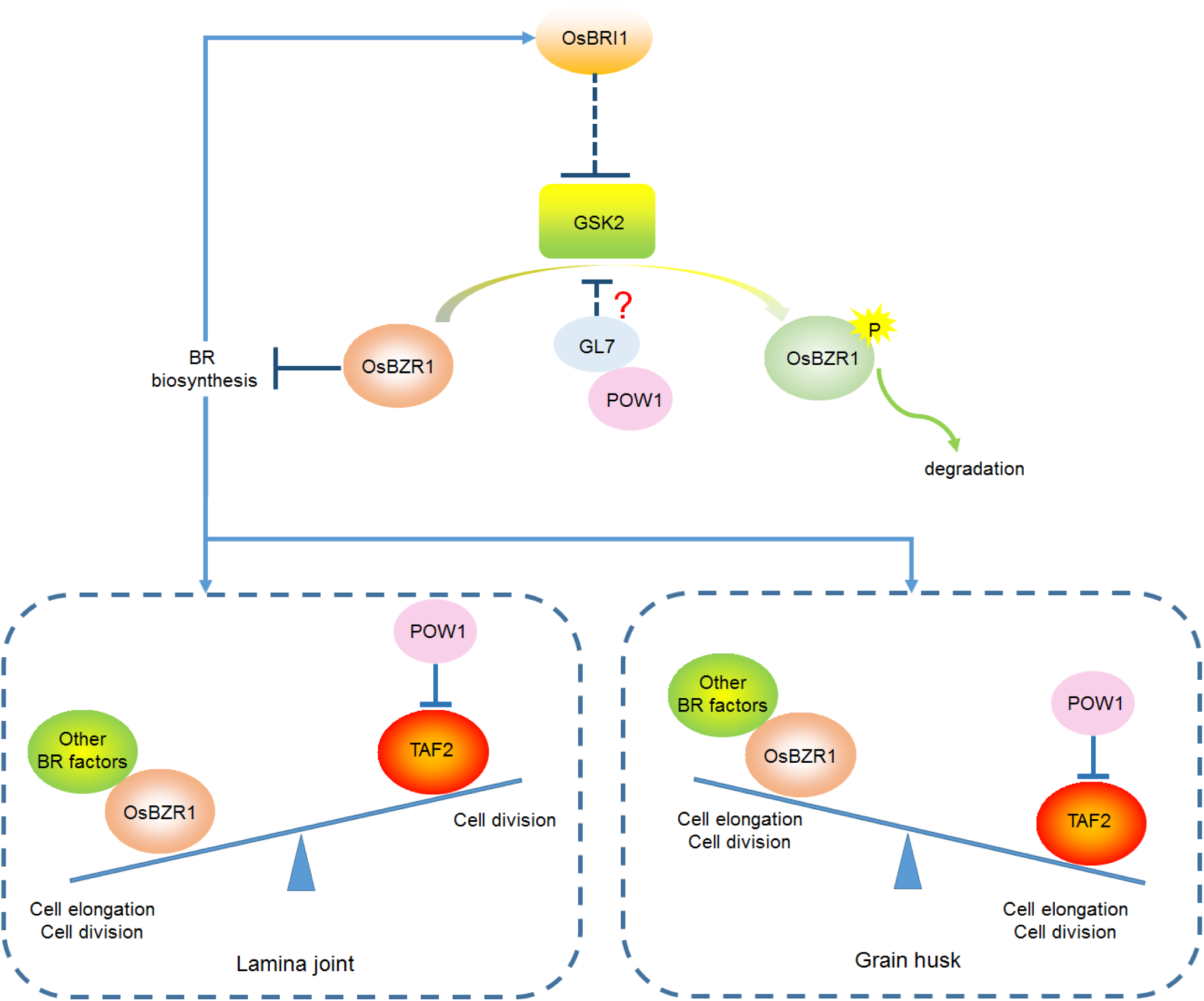
The putative working model of *POW1* in separable regulation of grain size and leaf angle development. POW1 is a key factor in the BR pathway by affecting OsBZR1 phosphorylation, and participates in *TAF2*-mediated cell cycle progression through inhibiting TAF2 transactivation activity. In the lamina joint, the little effect of *TAF2* on cell elongation leads to the dominant role of BR in leaf angle formation. While in the young panicle, the marked role of *TAF2* in cell division and cell elongation gives rise to the TAF2-dominated grain size development. The red question mark indicates that it is not clear whether GL7 is the direct linker between POW1 and GSK2 to regulate the phosphorylation of OsBZR1.

### The cellular processes in certain tissues are variably determined by the equilibrium among *POW1-TAF2-OsBZR1*

As a well-studied phytohormone, BR is involved in both leaf angle formation and grain size regulation (Tong and Chu, 2018), and one of the most amazing phenomena revealed in this study is that BR only affects the development of leaf angle but not grain size under the *pow1* mutant background. This observation might be explained by that BR shows differential roles in certain tissues (Tong and Chu, 2018), and lamina joint is one of the most sensitive tissues responding to BR fluctuation (Morinaka et al., 2006). Therefore, the effect caused by reduction or increment in endogenous BR levels would be more prominent for the development of leaf angle than that of grain size. Moreover, *OsBZR1*, the central factor of the BR pathway, was reported to govern cell elongation (Tong et al., 2014; Tong and Chu, 2018), and adaxial cell expansion was proven to be mainly responsible for the BR-induced leaf angle formation (Cao and Chen, 1995). Accordingly, the enlarged leaf angle phenotype of *pow1* could be readily rescued by manipulation of the BR pathway components. In contrast to the effect of BR on cell elongation, *TAF2* shows little effects on cell expansion during leaf angle formation, and expression of this gene is not strongly induced in the lamina joint of the *pow1* mutant. However, transcription of *TAF2* is sharply induced in the young panicle of *pow1*, resulting in drastic change in grain epidermal cell size and cell number. Although dysfunction of genes involved in BR biosynthesis and signaling could effectively modify grain development under the WT background, the grain size of *pow1^D4-D11^*^−RNAi^ and *pow1/d61* plants remained unchangeable compared to that of the mutant. These results suggest that the BR pathway functions upstream of the *POW1*-*TAF2*-mediated cellular processes, and manipulation of the BR pathway under the *pow1* background therefore could not antagonize the enhanced transcription and transactivation activity of the downstream *TAF2* during grain size formation. We suppose that the separable functions of *POW1* in the regulation of leaf angle and grain size might be attributed to that in these two tissues, *TAF2* shows differential responses to the *pow1* mutation at the transcription level and distinct roles in cell elongation and cell division. In the lamina joint, the less induction of *TAF2* expression and the little effect of this gene on cell elongation lead to the dominant role of BR in leaf angle formation under the mutant background. Contrastingly, in the young panicle, the drastic increase of transcription and transactivation activity of TAF2, and the significant effect of *TAF2* on epidermal cell size and number give rise to the TAF2-dominated grain size development in *pow1*. In conclusion, our results indicate that POW1 is a key and global factor in the BR pathway by affecting OsBZR1 phosphorylation, and participates directly in *TAF2*-mediated cell cycle progression through influencing TAF2 transactivation activity, thus making the *pow1* phenotype difficult to be rescued by any one of the two factors (Figure 11).

### *POW1* has great potential in high yield breeding

The erect-leaf trait is considered to be the ideotype for photosynthesis, growth and grain production (Sakamoto et al., 2006), and has attracted wide attention in especially molecular design of crop varieties with high yield potential. In rice, genetic analysis has revealed that overexpression of *OsAGO7* could decrease leaf angle by inducing upward leaf curling. However, this gene also shows adverse effects on other traits such as chlorophyll content (Shi et al., 2007). Loss of function of *OsILA1* could increase leaf angle by altering vascular bundle formation and cell wall composition. However, there is no evidence that genetic manipulation of this gene could make the leaf erect (Ning et al., 2011). Many studies have shown that leaf angle development is closely related to plant hormones including auxin (Cao and Chen, 1995), BR (Tong and Chu, 2018), and gibberellin (Shimada et al., 2006), and these findings have provided voluable information for understanding the molecular mechanism controlling leaf angle formation. However, although genetic manipulation of the related genes could effectively decrease the leaf angle, many other traits such as plant height and fertility would be adversely modified. Especially, the development of leaf angle is in most cases directly or indirectly related to BR (Tong and Chu, 2018), which also affects the development of grain size, another key yield determinant. Although the decrease of leaf angle by modifying BR pathway is often accompanied by the reduction of grain size, Morinaka et al., (2006) found that moderate suppression of *OsBRI1* expression could yield plants with erect-leaf phenotype without grain changes. In addition, loss of function mutant of *D4* displayed slight dwarfism and erect leaves without undesirable phenotypes such as reduction in grain size (Sakamoto et al., 2006). With standard fertilizer application, the grain yield of the *d4* mutant under dense planting increased substantially compared to that of the WT under conventional conditions. In this study, we show that *pow1* shows typical BR-related phenotypes of enlarged grain size and leaf angle. However, unlike that regulated by BR pathway genes, these two traits in *pow1* could be separately controlled by modulating the expression of *TAF2* and *D4/D11*, respectively. The separable regulation of *POW1* in grain size and leaf angle development thus provides a promising strategy to design high-yielding varieties with not only compact plant architecture but also increased grain size, as observed in the *pow1^D4-D11-^*^RNAi^ transgenic plants and the *pow1/d61* double mutant plants. In other words, by suppressing the expression of *TAF2* under the *pow1^D4-D11-^*^RNAi^ background, the grain size could be freely modified without altering the erect-leaf phenotype. Therefore, compared with the previous findings (Morinaka et al., 2006; Sakamoto et al., 2006), our results described here suggest that utilization of the *POW1-TAF2-*BR pathway could promote high yield breeding a further step forward in rice.

## METHODS

### Plant materials and growth conditions

The *pow1* mutant was isolated from the M_2_ population of the *japonica* cultivar KY131 mutated with sodium azide. The mapping population was generated by crossing *pow1* with the *indica* cultivar Kasalath. To create the double mutants *pow1/d61-1* and *pow1/d61-2, pow1* was used as the female to cross with *d61-1* and *d61-2* mutants, and the resulting F_1_ plants were backcrossed with *pow1* three times, respectively. The self-fertilized BC_3_F_2_ plants were genotyped with gene specific markers to select the double plants. To exclude background noise, we also selected all single mutants from the same genetic population for phenotypic comparison. All plants used in this study were cultivated in the experimental fields in Beijing or Sanya (Hainan Province) during the natural growing season. The primers used for mutant selection are listed in Supplemental Table 1.

### Lamina joint assay and BL and BRZ treatments

For the lamina joint assay, the second leaf laminas of 7-day-old seedlings grown in the dark were excised and submerged in distilled water that contained different concentrations of BL (Sigma), and phenotypic analysis was performed after 3 days of treatment. To observe coleoptile elongation, the seeds were sterilized with 3% NaClO, sowed on half strength solid MS medium that contained different concentrations of BL, and grew for 3 additional days after germination. For BRZ treatment, the sterilized seeds were sowed on half strength MS medium that contained 0, 5 and 20 μM BRZ and grew for one week after germination. For the BR induction assay, two-week-old seedlings cultured in a greenhouse were transplanted into half strength MS liquid medium that contained 1 μM BL, and shoots were sampled after 0, 2, 6, 12, and 24 h treatment for qRT-PCR assay.

### Histological analysis

To prepare paraffin sections, samples were fixed in FAA solution (50% ethanol, 5% acetic acid and 3.7% formaldehyde) overnight at 4°C after 15 min vacuum treatment; the samples were subsequently dehydrated with a graded ethanol series and embedded in paraplast for 2 days at 60°C. Sections (8 μm) were prepared with a microtome (RM2235, Leica), stained with 0.5% toluidine blue and observed with a microscope (BX53, Olympus).

For SEM analysis, glumes were collected prior to grain filling and fixed in 1× PBS solution that contained 2.5% glutaraldehyde overnight at 4°C. After dehydration with a series of ethanol and substitution with ethyl acetate, the samples were critical-point dried, sprayed with gold particles and observed with a scanning electron microscope (S-3000N, Hitachi). Cells of the middle part of the lemma were observed for imaging and entire grain were imaged for cell number counting.

### Quantification of endogenous BR content

Fresh leaves of two-month-old plants of *pow1* and WT were frozen in liquid nitrogen and then grounded to a fine powder. BR quantification was performed based on a previously described method (Xin et al., 2013).

### Map-based cloning

The mapping population was constructed by crossing *pow1* with the *indica* cultivar Kasalath. Using 20 bulked WT and mutant plants, respectively, the candidate gene was first mapped to the short arm of chromosome 7 between the SSR markers RM8010 and RM1353. Further analysis of the F_2_ mutant plants subsequently fine-mapped the casual gene to the region between RM21072 and RM21057, and sequence comparisons were then performed between *pow1* and WT for all annotated genes within this region.

### qRT-PCR assay

Total RNA was extracted using the RNAios PLUS reagent (Takara), and 1 μg of RNA was reverse-transcribed by the oligo (dT) primer using a reverse transcription kit (Promega) after digestion with RNase-free DNaseI (Fermentas). The qRT-PCR assay was performed in triplicate with SYBR Green I Master reagent and the Light Cycler Nano system (Roche). *Actin* was used as the internal control for normalization. The primers used in this study are listed in Supplemental Table 1.

### Vector construction and plant transformation

For the complementation test, a 4.6-kb genomic fragment that contained the entire wild type *POW1* genomic sequence, including the 2.2-kb native promoter, was cloned into the pZH2B vector. To create *D4, D11* and *TAF2* single RNAi plants, the hairpin sequence with two ∼300-bp cDNA inverted repeats was inserted into pZH2Bi. For *D4* and *D11* double RNAi construct, the cassette, including the ubiquitin promoter, hairpin sequence and Nos terminator, was cut from the *D4*-pZH2Bi construct and inserted into the *D11*-pZH2Bi construct. These vectors were transformed into WT or *pow1* with the *Agrobacterium tumefaciens*-mediated transformation method. The primers used for vector construction are listed in Supplemental Table 1.

### Transient expression assay in rice protoplast

For subcellular localization, the full-length coding sequence of *POW1/OsBZR1* was cloned into the pSAT6-EYFP-N1 vector to form the *POW1/OsBZR1-EYFP* (*Enhanced yellow fluorescent protein*) construct. Empty vector was used as a negative control. For the BiFC assay, the coding sequences of *POW1*, *OsBZR1* and *TAF2*-C were PCR amplified, inserted into the pUC19-VYNE (R) or pUC19-VYCE (R) vector and fused with the N- or C-terminus of the Venus YFP sequence, respectively. These vectors and the corresponding empty vectors were cotransformed in different combinations into rice protoplast. The YFP signal was detected with a confocal laser scanning microscope (CLSM; FV1000, Olympus) after 16 h incubation at 28°C in the dark. For the transactivation assay, the coding sequence of *POW1* was amplified and fused with GAL4 DNA binding domain in the pRT-BD vector, and the resulting construct was used as the effectors. The LUC vector, which contains the GAL4 binding motif and luciferase coding region, was used as the reporter, and the vector that expressed *Renilla* luciferase (pTRL) was employed as the internal control. The effector was then cotransformed into rice protoplast with the reporter and the internal control, respectively.

To detect the transactivation activity of TAF2 on cell division and expansion genes, the coding sequences of *TAF2* were PCR amplified and inserted into pSAT6-EYFP-N1, thus the TAF2-YFP was used as the effector. The *35S* promoter in LUC was replaced by the promoters of cell division and expansion genes (∼2.0-kb) and the resulting constructs were used as reporters. The pTRL vector was then cotransformed with the resulting constructs into rice protoplast. The empty pSAT6-EYFP-N1 vector was also cotransformed as a negative control.

To detect the repressive effects of POW1 on TAF2, the coding sequences of *POW1*, *mPOW1* were PCR amplified and inserted into pSAT6-EYFP-N1, respectively. These two constructs were used as co-effectors with TAF2-YFP.

To detect OsBZR1-mediated inhibition on *D4* and *D11* expression, the promoters (∼2.0-kb) of *D4* and *D11* were cloned into the LUC vector by replacing the *35S* promoter, respectively, and the resulting constructs were used as reporters. To create the effector construct, the coding sequence of *OsBZR1* was cloned into pSAT6-EYFP-N1 vector. To diminish the background difference, the empty pSAT6-EYFP-N1 vector was used as the negative control to cotransform with each reporter construct into the protoplast of *pow1* and WT, respectively. The pTRL vector was also cotransfected as an internal control.

After 16 h incubation at 28°C in the dark, the protoplast was centrifuged at 150 × *g*, and the pellet was used for the dual-luciferase assay as described in the Dual-Luciferase® Reporter Assay System Manual (Promega). All transformations were performed via the PEG (polyethylene glycol)/CaCl_2_-mediated method. The primers used for the transient assay are listed in Supplemental Table 1.

### Yeast two-hybrid assay

The coding sequences of *POW1*, *TAF2*, *TAF2-*C and *OsBZR1* were cloned into the pGBKT7 or pGADT7 vector (Clontech), respectively, and the resulting constructs and the corresponding empty vectors were then transformed into the yeast strain Golden Yeast with different combinations. Interactions were detected on SD/-Leu-Trp-His-Ade medium or SD/-Leu-Trp-His medium. The transformation was performed as described in the Clontech Yeast Two-Hybrid System User Manual. The primers used for the yeast two-hybrid assay are listed in Supplemental Table 1.

### Protein sequence alignment

The amino acid sequence of POW1 was compared with the *Arabidopsis* homolog ALP1, the known DDE-domain containing transposonase Hs Harbi1 from human (*Homo sapiens*), and Dr Harbi1 and XP_009300611.1 from zebrafish (*Danio rerio*). Alignment was performed using ClustalW. The sequences used for alignment were obtained by blasting with the POW1 protein sequence on the NCBI website (https://blast.ncbi.nlm.nih.gov/Blast.cgi).

### ChIP assay

Formaldehyde cross-linked chromatin DNA was isolated from two weeks old leaves of WT and *pow1*. Immuno-precipitation was performed by anti-H3K27me^3^ antibody (Millipore 07-449), then the H3K27me3 bounded DNA was isolated by protein A Dynabeads (Invitrogen, 10002D). The ChIP DNA was then used as a template for qRT-PCR assay. The input DNA before immunoprecipitation was used as control. Primers used are listed in Supplemental Table 1.

### *In vitro* and semi-*in vivo* pull down assay

GST-tagged protein and bounded by Glutathione beads (GE healthcare). 6×His-tagged protein were expressed and purified by Ni-NTA His Bind Resin (Millipore 70666-3). ∼0.2 μg of GST-OsBZR1, GST-POW1 and GST-mPOW1 bounded to GST beads were incubated with ∼0.3 μg TAF2-C-His at 4°C overnight, then the GST beads were collected by centrifugation and washed with 1×PBS for 5 times. The output protein was detected using anti-His antibody (Abmart M20020). For the detection of interaction between GL7 and POW1/GSK2, ∼0.2 μg of GST-POW1 or GST-GSK2 bounded beads were made to pull down GL7-YFP expressed in rice protoplast. The GST beads were then washed with protein extraction buffer (50 mM sodium phosphate buffer, pH 7.4, 150mM NaCl, 10% glycerol, 0.1% NP-40, 1× protease inhibitor cocktail) for 3 times. The output protein was detected using anti-GFP antibody (Abcam ab290).

### Phosphorylation analysis of OsBZR1 in plants

Flag leaves from two-month old plants were ground into powder in liquid and boiled with SDS-PAGE sample buffer. Protein samples were resolved by SDS-PAGE with or without 50 μM phos-tag (ApexBio F4002). OsBZR1 was detected by the commercial anti-OsBZR1 and anti-HSP was used as equal loading control (BPI, http://www.proteomics.org.cn).

### Accession numbers

Sequence data from this article could be found in the EMBL/GenBank data libraries under the following accession numbers: *POW1*, *Os07g07880*; *OsBZR1*, *Os07g39220*; *GSK2*, *Os05g11730*; *GL7*, *Os07g41200*; *ILI1*, *Os04g54900*; *RAVL1*, *Os04g49230*; *D2*, *Os01g10040*; *D4*, *Os03g12660*; *D11*, *Os04g39430*; *BRD1*, *Os03g40540*; *OsBRI1*, *Os01g52050* and *TAF2*, *Os09g24440*.

## Supplemental data

**Supplemental Figure 1.** Organ size comparison between *pow1* and WT. Scale bars, 2 cm for **(A)**, 5 cm for **(B)**, 2 cm for **(C)**, 2 mm for **(D)** and **(E)**, and 500 μm for **(F)**, respectively.

**Supplemental Figure 2.** Comparison of the inner epidermal cells of the lemma between *pow1* and WT, which indicates the cells in *pow1* were longer and wider than those in WT. Scale bars, 50 μm.

**Supplemental Figure 3.** Histological analysis of lamina joint of *pow1* and WT. **(A)** and **(C)** represent sections before the leaf angle forms, and **(B)** and **(D)** after the leaf angle forms. Red double-headed arrows in **(A)** and **(B)** indicate the adaxial cell of lamina joint. Scale bars, 100 μm. The left section in **(C)** and **(D)** show the adaxial side of lamina joint.

**Supplemental Figure 4.** Protein sequence alignment of POW1, ALP1 and HARBI1 proteins from human and zebrafish. Amino acids of ALP1 and POW1 are colored according to their properties. Black boxes indicate conserved DDE triads that are disrupted in both ALP1 and POW1.

**Supplemental Figure 5.** POW1 does not appear to function as an epigenetic regulator or a transcription factor.

**(A)** and **(B)** Transactivation activity assay. No significant transactivation activity was observed for POW1 in both yeast and rice protoplast.

**(C)** DNA methylation assay. No obvious difference of H3K27me3 levels within *D4* and *D11* gene locus was observed between *pow1* and WT. Data are means ± SE (*n* = 3). *P* values from the student’s *t*-test of *pow1* against WT were indicated.

**Supplemental Figure 6.** Downregulation of *D4* and *D11* shows little effects on grain size modification under the *pow1* mutant background.

**(A)** Expression analysis of *D4* and *D11* in the single and double RNAi transgenic plants. The transcript levels were normalized against WT, which was set to 1. Data are means ± SE (*n* = 3). Bars followed by the different letters represent significant difference at 5%.

**(B)** Grain length comparison among the *D4/D11* RNAi transgenic plants. Data are means ± SE (*n* = 15). Bars followed by the different letters represent significant difference at 5%.

**Supplemental Figure 7.** *pow1* shows enhanced BR signaling.

**(A)** Lamina joint assay. The second lamina joints of 7-day-old seedlings grown in the dark were excised and submerged in distilled water that contained different BL concentrations for 3 days. Data are means ± SE (*n* = 20). *P* values from the student’s *t*-test of the *pow1* against WT were indicated.

**(B)** Coleoptile elongation assay. The coleoptile length of *pow1* and WT grown for 3 days under different BL concentrations was compared. Data are means ± SE (*n* = 10). *P* values from the student’s *t*-test of the *pow1* against WT were indicated.

**(C)** Expression analysis of the BR signaling genes *OsBZR1, RAVL1, OsBRI1* and *ILI1*. The transcript levels were normalized against WT, which was set to 1. Data are means ± SE (*n* = 3). *P* values from the student’s *t*-test of the *pow1* against WT were indicated.

**Supplemental Figure 8.** Comparison of leaf angle and grain length among WT, *pow1*, *d61-1*, *d61-2*, and the double mutants. Data are means ± SE (*n* = 20). Bars followed by the different letters represent significant difference at 5%.

**Supplemental Figure 9.** Cytological observation. SEM observation and statistical analysis were conducted for the outer epidermal cells of grain husks from WT and *POW1^TAF2^*^−RNAi^ plants (*n* = 20 for cell size and 5 for cell number). *P* values from the student’s *t*-test were indicated. Scale bars, 100 μm.

**Supplemental Figure 10.** Sketch of the constructs used for luciferase assay.

**(A)** Constructs for detecting TAF2 transactivation activity on cell cycle and expansion genes.

**(B)** Constructs for detecting the effect of POW1/mPOW1 on the transactivation activity of TAF2 on cell cycle and expansion genes.

**(C)** Constructs for detecting the inhibitive activity of OsBZR1 on *D4* and *D11* expression.

**Supplemental Figure 11.** Histological analysis of lamina joint. (**C)**, (**D)**, (**G)**, and (**H)** indicate the adaxial side of lamina joint. Scale bars, 50 μm.

**Supplemental Figure 12.** Interaction assay of POW1-OsBZR1 and TAF2-OsBZR1. No direct interaction was detected between POW1/TAF2-C and OsBZR1 in both yeast **(A)** and rice protoplast **(B)**.

**Supplemental Table 1.** Primers used in this study.

## ACKNOWLEDGEMENTS

This work was supported by grants from the National Key Research and Development Program of China (Grant: 2016YFD0101801) and the State Key Laboratory of Plant Genomics (SKLPG2011B0403). S.F. and J.C. are supported by the National Natural Science Foundation of China (3147043). We thank Professor Hongning Tong (Chinese Academy of Agricultural Sciences) for suggestions regarding exogenous BRZ treatments. The *d61* mutants were kindly provided by Professor Chengcai Chu, and the pRT-BD, pTRL and LUC vectors were provided by Professor Shouyi Chen (Institute of Genetics and Developmental Biology, Chinese Academy of Sciences).

## AUTHOR CONTRIBUTIONS

S.Y. conceived and supervised the project. S.Y. and L.Z. designed the study and analyzed the data. L.Z. performed the functional analyses. R.W. screened the *pow1* mutants, created mapping population and double mutants, and carried out field management. Y.W. contributed the reagents and equipment management. S.F. and J.C. performed the BR quantification. Y.X. and L.Z. prepared the photos. S.Y. and L.Z. wrote the manuscript. All authors have read and approved the final manuscript.

## REFERENCES

Asahara, H., Santoso, B., Guzman, E., Du, K., Cole, P.A., Davidson, I., and Montminy, M. (2001). Chromatin-dependent cooperativity between constitutive and inducible activation domains in CREB. Mol. Cell. Biol. 21, 7892–7900.

Asami, T., Mizutani, M., Fujioka, S., Goda, H., Min, Y.K., Shimada, Y., Nakano, T., Takatsuto, S., Matsuyama, T., Nagata, N., Sakata, K., and Yoshida, S. (2001). Selective interaction of triazole derivatives with DWF4, a cytochrome P450 monooxygenase of the brassinosteroid biosynthetic pathway, correlates with brassinosteroid deficiency in *Planta*. J. Biol. Chem. 276, 25687–25691.

Bahat, A., Kedmi, R., Gazit, K., Richardo-Lax, I., Ainbinder, E., and Dikstein, R. (2013). TAF4b and TAF4 differentially regulate mouse embryonic stem cells maintenance and proliferation. Genes Cells 18, 225–237.

Bai, M.Y., Zhang, L.Y., Gampala, S.S., Zhu, S.W., Song, W.Y., Chong, K., and Wang, Z.Y. (2007). Functions of OsBZR1 and 14-3-3 proteins in brassinosteroid signaling in rice. Proc. Natl. Acad. Sci. USA 104, 13839–13844.

Bertrand, C., Benhamed, M., Li, Y.F., Ayadi, M., Lemonnier, G., Renou, J.P., Delarue, M., and Zhou, D.X. (2005). Arabidopsis HAF2 gene encoding TATA-binding protein (TBP)-associated factor TAF1, is required to integrate light signals to regulate gene expression and growth. J. Biol. Chem. 280, 1465–1473.

Cao, H., and Chen, S. (1995). Brassinosteroid-induced rice lamina joint inclination and its relation to indole-3-acetic acid and ethylene. Plant Growth Regul. 16, 189–196.

Chen, Z., and Manley, J.L. (2000). Robust mRNA transcription in chicken DT40 cells depleted of TAFII31 suggests both functional degeneracy and evolutionary divergence. Mol. Cell. Biol. 20, 5064–5076.

Duan, Y., Li, S., Chen, Z., Zheng, L., Diao, Z., Zhou, Y., Lan, T., Guan, H., Pan, R., Xue, Y., and Wu, W. (2012). *Dwarf and deformed flower 1*, encoding an F-box protein, is critical for vegetative and floral development in rice (*Oryza sativa* L.). Plant J. 72, 829–842.

Eom, H., Park, S.J., Kim, M.K., Kim, H., Kang, H., and Lee, I. (2018). TAF15b, involved in the autonomous pathway for flowering, represses transcription of FLOWERING LOCUS C. Plant J. 93, 79–91.

Garbett, K.A., Tripathi, M.K., Cencki, B., Layer, J.H., and Weil, P.A. (2007). Yeast TFIID serves as a coactivator for Rap1p by direct protein-protein interaction. Mol. Cell. Biol. 27, 297–311.

Gupta, K., Sari-Ak, D., Haffke, M., Trowitzsch, S., and Berger, I. (2016). Zooming in on transcription preinitiation. J. Mol. Biol. 428, 2581–2591.

Hong, Z., Ueguchi Tanaka, M., Umemura, K., Uozu, S., Fujioka, S., Takatsuto, S., Yoshida, S., Ashikari, M., Kitano, H., and Matsuoka, M. (2003). A rice brassinosteroid-deficient mutant, *ebisu dwarf* (*d2*), is caused by a loss of function of a new member of cytochrome P450. Plant Cell 15, 2900–2910.

Jacobson, R.H., Ladurner, A.G., King, D.S., and Tjian, R. (2000). Structure and function of a Human TAFII 250 double bromodomain module. Science (New York, N.Y.) 288, 1422–1425.

Jiang, P., Wang, S., Zheng, H., Li, H., Zhang, F., Su, Y., Xu, Z., Lin, H., Qian, Q., and Ding, Y. (2018). SIP1 participates in regulation of flowering time in rice by recruiting OsTrx1 to *Ehd1*. New Phytol. 219, 422–435.

Jiang, Y., Bao, L., Jeong, S.Y., Kim, S.K., Xu, C., Li, X., and Zhang, Q. (2012). XIAO is involved in the control of organ size by contributing to the regulation of signaling and homeostasis of brassinosteroids and cell cycling in rice. Plant J. 70, 398–408.

Kapitonov, V.V., and Jurka, J. (1999). Molecular paleontology of transposable elements from *Arabidopsis thaliana*. Genetica 107, 27–37.

Kapitonov, V.V., and Jurka, J. (2004). Harbinger transposons and an ancient HARBI1 gene derived from a transposase. DNA Cell Biol. 23, 311–324.

Kim, B.K., Fujioka, S., Takatsuto, S., Tsujimoto, M., and Choe, S. (2008). Castasterone is a likely end product of brassinosteroid biosynthetic pathway in rice. Biochem. Biophys. Res. Commun. 374, 614–619.

Lago, C., Clerici, E., Mizzi, L., Colombo, L., and Kater, M.M. (2004). TBP-associated factors in Arabidopsis. Gene 342, 231–241.

Lago, C., Clerici, E., Dreni, L., Horlow, C., Caporali, E., Colombo, L., and Kater, M.M. (2005). The Arabidopsis TFIID factor AtTAF6 controls pollen tube growth. Dev. Biol. 285, 91–100.

Li, N., Xu, R., Duan, P., and Li, Y. (2018). Control of grain size in rice. In Plant Reproduction, pp. 1–15.

Liang, S.C., Hartwig, B., Perera, P., Mora-García, S., de Leau, E., Thornton, H., de Alves, F.L., Rapsilber, J., Yang, S., James, G.V., Schneeberger, K., Finnegan, E.J., Turck, F., and Goodrich, J. (2015). Kicking against the PRCs – A domesticated transposase antagonises silencing mediated by Polycomb Group Proteins and is an accessory component of polycomb repressive complex 2. PLoS Genet. 11, e1005660.

Lindner, M., Simonini, S., Kooiker, M., Gagliardini, V., Somssich, M., Hohenstatt, M., Simon, R., Grossniklaus, U., and Kater, M.M. (2013). TAF13 interacts with PRC2 members and is essential for Arabidopsis seed development. Dev. Biol. 379, 28–37.

Liu, J., Chen, J., Zheng, X., Wu, F., Lin, Q., Heng, Y., Tian, P., Cheng, Z., Yu, X., Zhou, K., Zhang, X., Guo, X., Wang, J., Wang, H., and Wan, J. (2017). GW5 acts in the brassinosteroid signalling pathway to regulate grain width and weight in rice. Nat. Plants 3.

Metzger, D., Scheer, E., Soldatov, A., and Tora, L. (1999). Mammalian TAFII30 is required for cell cycle progression and specific cellular differentiation programmes. EMBO J. 18, 4823–4834.

Mori, M., Nomura, T., Ooka, H., Ishizaka, M., Yokota, T., Sugimoto, K., Okabe, K., Kajiwara, H., Satoh, K., Yamamoto, K., Hirochika, H., and Kikuchi, S. (2002). Isolation and characterization of a rice dwarf mutant with a defect in brassinosteroid biosynthesis. Plant Physiol. 130, 1152–1161.

Morinaka, Y., Sakamoto, T., Inukai, Y., Agetsuma, M., Kitano, H., Ashikari, M., and Matsuoka, M. (2006). Morphological alteration caused by brassinosteroid insensitivity increases the biomass and grain production of rice. Plant Physiol. 141, 924–931.

Ning, J., Zhang, B., Wang, N., Zhou, Y., and Xiong, L. (2011). Increased Leaf Angle1, a Raf-Like MAPKKK that interacts with a nuclear protein family, regulates mechanical tissue formation in the lamina joint of rice. Plant Cell 23, 4334–4347.

Nogales, E., Louder, R.K., and He, Y. (2017). Structural insights into the eukaryotic transcription initiation machinery. Annu. Rev. Biophys. 46, 59–83.

Qiao, S.L., Sun, S.Y., Wang, L.L., Wu, Z.H., Li, C.X., Li, X.M., Wang, T., Leng, L.N., Tian, W.S., Lu, T.G., and Wang, X.L. (2017). The RLA1/SMOS1 transcription factor functions with OsBZR1 to regulate brassinosteroid signaling and rice architecture. Plant Cell 29, 292–309.

Reeves, W.M., and Hahn, S. (2005). Targets of the Gal4 transcription activator in functional transcription complexes. Mol. Cell. Biol. 25, 9092–9102.

Rojo Niersbach, E., Furukawa, T., and Tanese, N. (1999). Genetic dissection of hTAFII130 defines a hydrophobic surface required for interaction with glutamine-rich activators. J. Biol. Chem. 274, 33778–33784.

Sakamoto, T., Morinaka, Y., Ohnishi, T., Sunohara, H., Fujioka, S., Ueguchi Tanaka, M., Mizutani, M., Sakata, K., Takatsuto, S., Yoshida, S., Tanaka, H., Kitano, H., and Matsuoka, M. (2006). Erect leaves caused by brassinosteroid deficiency increase biomass production and grain yield in rice. Nat. Biotech. 24, 105–109.

Sekiguchi, T., Nohiro, Y., Nakamura, Y., Hisamoto, N., and Nishimoto, T. (1991). The human CCG1 gene, essential for progression of the G1 phase, encodes a 210-kilodalton nuclear DNA-binding protein. Mol. Cell. Biol. 11, 3317–3325.

Shi, Z., Wang, J., Wan, X., Shen, G., Wang, X., and Zhang, J. (2007). Over-expression of rice *OsAGO7* gene induces upward curling of the leaf blade that enhanced erect-leaf habit. Planta 226, 99–108.

Shimada, A., Ueguchi-Tanaka, M., Sakamoto, T., Fujioka, S., Takatsuto, S., Yoshida, S., Sazuka, T., Ashikari, M., and Matsuoka, M. (2006). The rice SPINDLY gene functions as a negative regulator of gibberellin signaling by controlling the suppressive function of the DELLA protein, SLR1, and modulating brassinosteroid synthesis. Plant J. 48, 390–402.

Sridhar, V.V., Surendrarao, A., and Liu, Z. (2006). *APETALA1* and SEPALLATA3 interact with SEUSS to mediate transcription repression during flower development Development 133, 3159–3166.

Tamada, Y., Nakamori, K., Nakatani, H., Matsuda, K., Hata, S., Furumoto, T., and Izui, K. (2007). Temporary expression of the *TAF10* gene and its requirement for normal development of *Arabidopsis thaliana*. Plant Cell Physiol. 48, 134–146.

Tanabe, S., Ashikari, M., Fujioka, S., Takatsuto, S., Yoshida, S., Yano, M., Yoshimura, A., Kitano, H., Matsuoka, M., Fujisawa, Y., Kato, H., and Iwasaki, Y. (2005). A novel cytochrome P450 is implicated in brassinosteroid biosynthesis via the characterization of a rice dwarf mutant, *dwarf11*, with reduced seed length. Plant Cell 17, 776–790.

Tian, J., Wang, C., Xia, J., Wu, L., Xu, G., Wu, W., Li, D., Qin, W., Han, X., Chen, Q., Jin, W., and Tian, F. (2019). Teosinte ligule allele narrows plant architecture and enhances high-density maize yields. Science (New York, N.Y.) 365, 658–664.

Tong, H., and Chu, C. (2018). Functional specificities of brassinosteroid and potential utilization for crop improvement. Trends Plant Sci. 23, 1016–1028.

Tong, H., Liu, L., Jin, Y., Du, L., Yin, Y., Qian, Q., Zhu, L., and Chu, C. (2012). DWARF AND LOW-TILLERING acts as a direct downstream target of a GSK3/SHAGGY-Like kinase to mediate brassinosteroid responses in rice. Plant Cell 24, 2562–2577.

Tong, H., Jin, Y., Liu, W., Li, F., Fang, J., Yin, Y., Qian, Q., Zhu, L., and Chu, C. (2009). DWARF AND LOW-TILLERING, a new member of the GRAS family, plays positive roles in brassinosteroid signaling in rice. Plant J. 58, 803–816.

Tong, H., Xiao, Y., Liu, D., Gao, S., Liu, L., Yin, Y., Jin, Y., Qian, Q., and Chu, C. (2014). Brassinosteroid regulates cell elongation by modulating gibberellin metabolism in rice. Plant Cell 26, 4376–4393.

Wang, S.K., Li, S., Liu, Q., Wu, K., Zhang, J.Q., Wang, S.S., Wang, Y., Chen, X.B., Zhang, Y., Gao, C.X., Wang, F., Huang, H.X., and Fu, X.D. (2015a). The *OsSPL16-GW7* regulatory module determines grain shape and simultaneously improves rice yield and grain quality. Nat. Genet. 47, 949–954.

Wang, Y., Xiong, G., Hu, J., Jiang, L., Yu, H., Xu, J., Fang, Y., Zeng, L., Xu, E., Xu, J., Ye, W., Meng, X., Liu, R., Chen, H., Jing, Y., Wang, Y., Zhu, X., Li, J., and Qian, Q. (2015b). Copy number variation at the *GL7* locus contributes to grain size diversity in rice. Nat. Genet. 47, 944.

Wang, Z.Y., Bai, M.Y., Oh, E., and Zhu, J.Y. (2012). Brassinosteroid signaling network and regulation of photomorphogenesis. Annu. Rev. Genet. 46, 701–724.

Waterworth, W.M., Drury, G.E., Hunter, G.B., and West, C.E. (2015). Arabidopsis TAF1 is an MRE11-interacting protein required for resistance to genotoxic stress and viability of the male gametophyte. Plant J. 84, 545–557.

Weake, V.M., and Workman, J.L. (2010). Inducible gene expression: diverse regulatory mechanisms. Nat. Rev. Genet. 11, 426–437.

Wu, C.Y., Trieu, A., Radhakrishnan, P., Kwok, S.F., Harris, S., Zhang, K., Wang, J.L., Wan, J.M., Zhai, H.Q., Takatsuto, S., Matsumoto, S., Fujioka, S., Feldmann, K.A., and Pennell, R.I. (2008). Brassinosteroids regulate grain filling in rice. Plant Cell 20, 2130–2145.

Xin, P., Yan, J., Fan, J., Chu, J., and Yan, C. (2013). An improved simplified high-sensitivity quantification method for determining brassinosteroids in different tissues of rice and Arabidopsis. Plant Physiol. 162, 2056–2066.

Yamamuro, C., Ihara, Y., Wu, X., Noguchi, T., Fujioka, S., Takatsuto, S., Ashikari, M., Kitano, H., and Matsuoka, M. (2000). Loss of function of a rice *brassinosteroid insensitive 1* homolog prevents internode elongation and bending of the lamina joint. Plant Cell 12, 1591–1605.

Yuan, Y.W., and Wessler, S.R. (2011). The catalytic domain of all eukaryotic cut-and-paste transposase superfamilies. Proc. Natl. Acad. Sci. USA 108, 7884–7889.

Zhu, X., Liang, W., Cui, X., Chen, M., Yin, C., Luo, Z., Zhu, J., Lucas, W.J., Wang, Z., and Zhang, D. (2015). Brassinosteroids promote development of rice pollen grains and seeds by triggering expression of Carbon Starved Anther, a MYB domain protein. Plant J. 82, 570–581.

Zuo, J., and Li, J. (2014). Molecular genetic dissection of quantitative trait loci regulating rice grain size. Annu. Rev. Genet. 48, 99–118.

